# Bioinformatic and Experimental Characterization of the RBM15 RNA Binding Protein

**DOI:** 10.1101/2023.07.20.549950

**Authors:** Emma Bose, Caleb Mayes, Lance Ellis, Corrine Baker, Sofia Tambalotti, Bryan Eusse, Shengwei Xiong, Yaa Pokua Osei Sarpong, Marwan Shalaby, Lucas Barry, Frank Lewis, Johnson Joseph, Talaidh Isaacs, Derik McCarthy, Dana Katz, Jingyang Wang, Victoria Zirimu, Luis Vargas, Julian Von Hofe, Glen C Aguilar, Katherine Buchan, Lei Zheng, Gregory Wolfe, Alisha N Jones

## Abstract

The RNA binding motif 15 protein (RBM15) plays a critical role in post-transcriptional regulation. Its role in facilitating N6-methyladenosine (m6A) modification, specifically through guiding the writer complex (WTAP METTL13 METTL14) to DRACH sequence motifs, has been demonstrated for several classes of RNA, including long noncoding RNAs (lncRNAs). The structural mechanism that underlies how RBM15 interacts with RNA has yet to be elucidated. In this study, we mined and bioinformatically assessed publicly available genome-wide RNA 2D structural probing and RBP cross-linking and immunoprecipitation data to investigate how RBM15 interacts with RNA, with a focus on lncRNA transcripts. RBM15, which possesses three RNA recognition motifs (RRMs), primarily interacts with stem-loop structured RNA motifs. Structural modeling reveals RRMs 2 and 3 are coaxially stacked in solution; these two RRMs are responsible for driving RBM15’s interaction with RNA. We further demonstrate this experimentally with two RNA hairpins, revealing low micromolar binding affinities. Altogether, this work provides insight into the structural mechanism by which RBM15 interacts with RNAs to govern biological function.

## 1. Introduction

The RNA binding motif 15 (RBM15) protein, also known as OTT1, is a member of the split end (SPEN) family of proteins ^1,2^. It possesses three N-terminal RNA recognition motifs (RRMs) and a Spen Orthologue and Paralogue C-terminal (SPOC) domain (**Figure 1A**). While RBM15 regulates various cellular processes—including RNA splicing, nuclear export, hematopoietic cell homeostasis, and chromosome inactivation—it is primarily recognized for its role as an accessory protein in the Wilms’ tumor 1-associated protein (WTAP)/methyltransferase-like 3 (METTL3)/methyltransferase-like 14 (METTL14) (WMM) writer complex ^3–9^. The WMM complex is responsible for the N6 methylation (m6A) of adenosines nested within DRACH motifs (where D = G/A/U, R = G/A, and H = A/C/U), a critical modification that influences RNA stability, splicing, and gene expression ^6,10–13^. RBM15, and its paralogue RBM15B, facilitate this process by positioning the methyltransferase complex close to the adenosines targeted for modification; the SPOC domain of RBM15 binds to WTAP and METTL3, while the RRMs interact with the RNA substrate (**Figure 1B**) ^6,14,15^.

**Figure 1.**
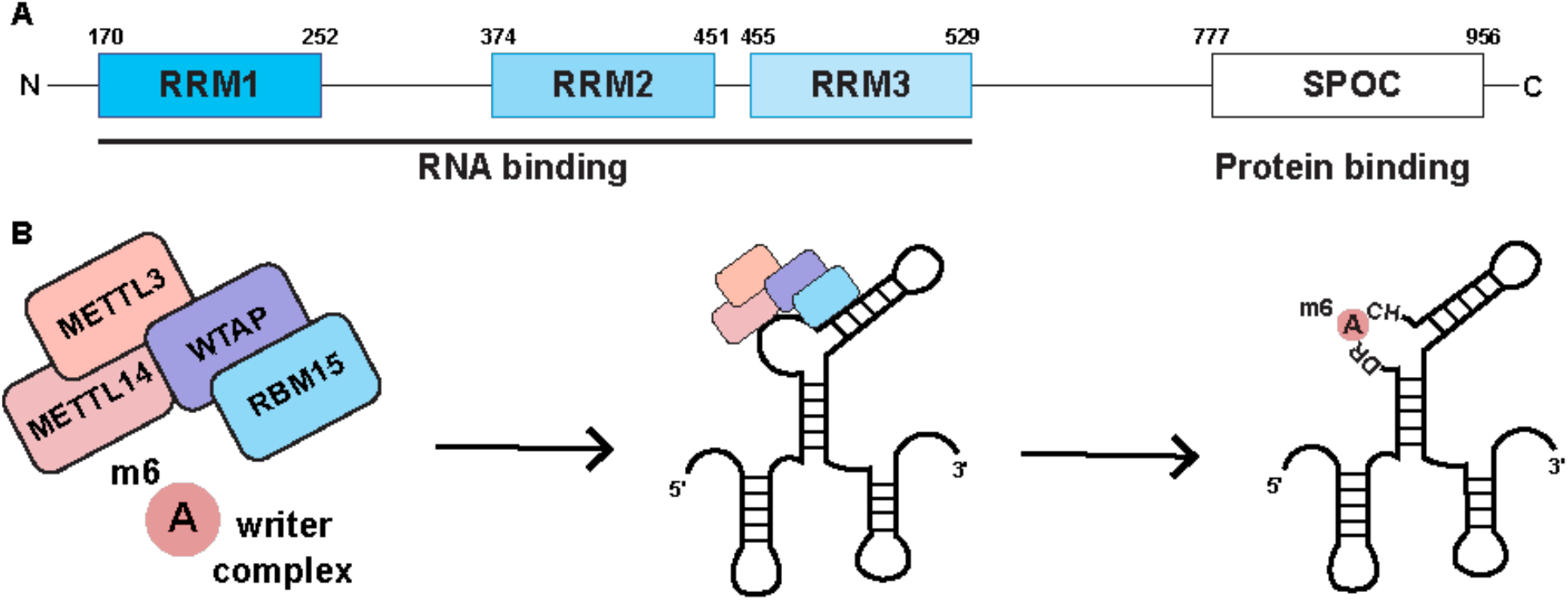
Schematic illustration of RBM15 and its functional role in m6A methylation. A) Protein domain architecture of RBM15, illustrating its key structural domains. B) Overview of RBM15’s role in guiding the m6A writer complex (WMM: WTAP, METTL3, and METTL14) to facilitate m6A modification (labeled in pink) within DRACH sequence motifs.

Several RNAs have been found to interact with RBM15, including two well-characterized long noncoding RNAs (lncRNAs): the X-inactive specific transcript (XIST) and the Functional Intergenic Repeating RNA Element (FIRRE) ^6,16–26^. XIST engages with nearly 80 RNA-binding proteins (RBPs) to transcriptionally silence the inactive X-chromosome in placental mammals ^26^. Silencing is facilitated by the A-repeats domain which is located at the 5’ end of the XIST transcript; this region is bound by RBM15 and undergoes m6A modification to regulate silencing ^6,27^. FIRRE, which is conserved across mammalian species, regulates numerous processes including adipogenesis, innate immunity, and hematopoiesis ^21–24,28^. It is composed of transposable elements known as local repeats (LRs) which facilitate interactions with nearly 70 RBPs, including RBM15 ^29^. Among the LRs is the RNA Repeating Domain (RRD) which is m6A modified, contributing to FIRRE’s functional roles in cells ^30^.

Though the roles of RBM15 in cells have been defined, the structural mechanism(s) by which RBM15 binds RNA have yet to be elucidated. In the absence of this information, we are limited in our ability to understand the structure-function relationship that underlies RBM15-RNA interactions. Here, we combine bioinformatic analyses and experimental assays to explore the structural mechanisms by which RBM15 binds to RNA. Using lncRNAs as a model system, we find i) RBM15 binds stem-loop structured motifs near DRACH motifs, and ii) RBM15 RRMs 2 and 3 form a heterodimer that drives its interaction with RNA.

## 2. Materials and Methods

### 2D structure prediction of lncRNAs

The LNCipedia was used to curate a list of all X-chromosome lncRNAs known to date. Genome-wide chemical probing datasets were downloaded from the RASP database ^31,51^. An in-house generated script was used to identify lncRNAs possessing chemical probing data in the RASP database and is available at: https://github.com/gpwolfe/match_lncRNA_chromosome.

The Ensemble project (release 109) was used to identify the Homo sapiens Ensemble Canonical transcript for each of the nine RNAs ^52^. Chemical probing data for the canonical transcript was mined from the RASP database. Using RNAStructure the secondary structure of each RNA was predicted (using chemical probing data as soft constraints) and the lowest energy structure (of twenty) was visualized using RNAcanvas ^53,54^.

### Identification of RNA-Protein binding partners and sites

Protein binding partners for each lncRNA were identified with oRNAment, and the binding with RBM15 was confirmed using CLIP data from ENCODE and POSTAR3 (**Table S1, Figure S1**) ^55–58^. We downloaded call sets for RBM15 eCLIP from the ENCODE portal with the following identifiers: ENCFF086HIT and ENCFF739LLZ ^59^.

CLIP (Crosslinking and Immunoprecipitation) data involves UV crosslinking of RNA-binding proteins (RBPs) to RNA molecules, followed by immunoprecipitation of the protein-RNA complexes and high-throughput sequencing of the RNA fragments that are crosslinked to the protein ^60,61^. In this study, CLIP data specifically targeting RBM15 were obtained from HEK293T and K562 cell lines to identify RBM15 interactions within each cellular context. It provides a comprehensive map of RBM15 binding sites on lncRNAs by pinpointing RNA regions that interact directly with this protein. Analysis of these binding sites enables the identification of sequence motifs or structural elements that may mediate protein-RNA interactions.

MEME (Multiple Em for Motif Elicitation) analysis was conducted on a dataset of 83 lncRNA sequences, shown to interact with RBM15 based on CLIP data, to identify motifs associated with RNA-binding protein (RBP) interactions ^62^. The Sensitive, Thorough, Rapid, Enriched Motif Elicitation (STREME) package within the MEME suite, optimized for the number of sequences over 50, was applied to facilitate this analysis ^63^. Sequences were retrieved from the BSgenome database and formatted in FASTA format for compatibility with MEME.

For motif discovery, a motif length threshold of 3 to 10 nucleotides was set, consistent with experimental data indicating that RBPs typically recognize sequences approximately 5-7 nucleotides in length. The results featured certain dinucleotides appearing as large letters in the motif output. These ‘big letters’ signify nucleotide positions with high frequency across the analyzed sequences, indicating potential key residues for binding affinity. This analysis identified statistically significant motifs enriched in RBM15 binding regions, suggesting conserved sequence patterns that may mediate RNA-protein interactions, particularly within the loop regions of stem-loop structures identified through CLIP data.

To identify the binding sites of these potential protein binders, catRAPID was used. Default settings were used for each query.

### *In vitro* transcription and purification of RNAs

The stem-loop RNAs from the XIST A-repeats and FIRRE RRD were *in vitro* transcribed using in-house prepared T7 polymerase. First, equal molar amounts of ssDNA template (with the T7 promoter) and T7 promoter-containing oligonucleotide were annealed by snap-cooling. Roughly 1 µM of these DNA templates was combined in 50 µL transcription reactions containing 20 mM MgCl_2_, 8 mM of each ribonucleoside triphosphate (rNTP), 5% polyethylene glycol 8000, 1X transcription buffer (5 mM Tris pH 8, 5 mM spermidine, 10 mM dithiothreitol (DTT)), 20% (v/v) dimethyl sulfoxide (DMSO) and 0.6 mg of T7 polymerase, which were then incubated at 37°C for 2 hours. Following transcription, the RNAs were purified on 20% urea denaturing polyacrylamide gels, followed by extraction of the RNA from the gel using two methods: electro-elution and crush and soak ^64^. Following, RNAs were ethanol precipitated in 1/10 volume of 3M sodium acetate and 3x total volume 100% ethanol, spun down at 13000 RPM, 4°C, for 1 hr; pellets were redissolved in nuclease-free water.

Table of RNA constructs:

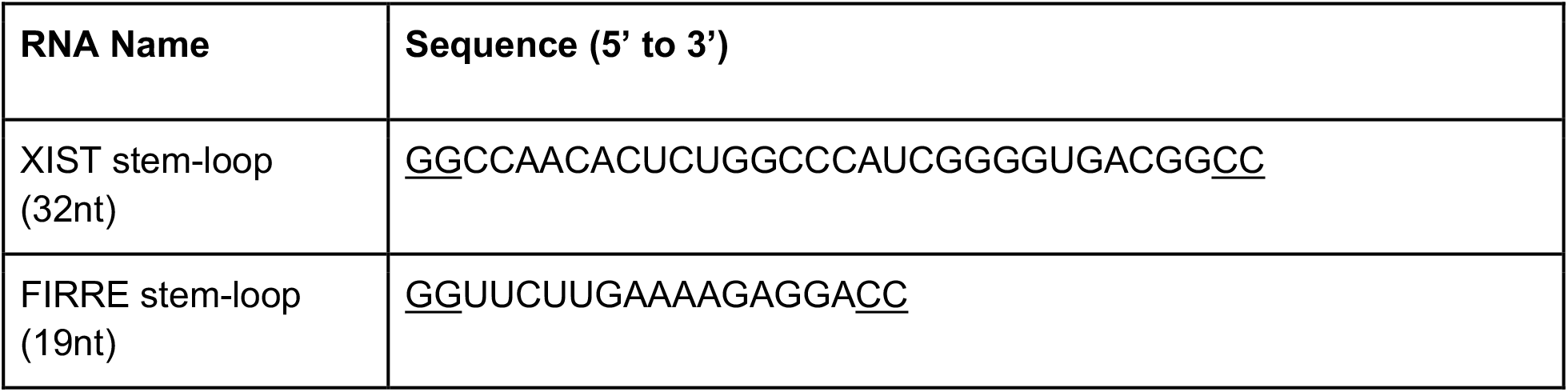

Underlined nucleotides indicate those added for transcriptional efficiency and to close the stem-loops.

### Protein expression and purification

DNA plasmid encoding each RBM15 construct was transformed into BL21 DE3 *E. coli* competent cells. Colonies were grown up in 1 L of lysogeny broth at 37°C to an optical density (OD_600_) of 0.9, and induced with 1 mM isopropyl beta-D-1-thiogalactopyranoside (IPTG) at 18°C for 16 hours. Cells were lysed by sonication in a buffer containing 300 mM NaCl, 10 mM imidazole, 50 mM Tris-HCl (pH 7.5), 5 mM beta-mercaptoethanol (BME), 1 mM 4-(2-aminoethyl) benzene sulfonyl fluoride hydrochloride (AEBSF).HCl, 1 mg/mL lysozyme, and 1 mg/mL deoxyribonuclease 1 (DNase1). The protein (possessing an N-terminal His_6_-tag followed by a Tobacco Etch Virus (TEV) protease cleavage site) was purified by immobilized metal affinity chromatography (IMAC) against an increasing imidazole concentration gradient. The N-terminal His_6_-tag was cleaved using in-house-prepared TEV protease, followed by reverse IMAC. The protein was further purified by size exclusion chromatography on a Superdex S75 purification column and eluted in a buffer containing 100 mM NaCl, 25 mM sodium phosphate pH 7.5, and 5 mM BME. Fractions containing purified protein were evaluated by sodium dodecyl-sulfate polyacrylamide gel electrophoresis (SDS PAGE), flash frozen in liquid nitrogen, and stored at −80°C in buffer containing 100 mM NaCl, 25 mM sodium phosphate pH 7.5, and 5 mM BME until further use.

### Filter dot blot assays

RNAs were 3’ labeled with Cy5 following established protocols ^65,66^. Following labeling, the RNAs were diluted to 50 nM in a reaction buffer containing 50 mM NaCl, 25 mM sodium phosphate (pH 7.5), and 5 mM BME. They were then mixed with increasing concentrations of protein as specified in the figures. The reactions were incubated at room temperature for 20 minutes to ensure equilibrium was reached before conducting the filter dot blot assays ^67^.

The filter dot blot was performed following established protocols ^41,68^. In brief, The filter dot blot was performed using a “dot-blot” apparatus (96-well BioRad Bio-Dot) with a double membrane setup: a 0.45 µm pore size positively charged nylon membrane (Whatman Nytran™SuperCharge) that retained free RNA and a 0.45 µm pore size nitrocellulose membrane (BioRad) that captured bound RNA. Before use, both membranes were presoaked for at least 30 min in a reaction buffer (50 mM NaCl, 25 mM sodium phosphate (pH 7.5), and 5 mM BME) to ensure optimal binding conditions. The samples were brought up to a final volume of 80 µL and then applied onto the double membrane setup, utilizing a vacuum pressure pump (StonyLab), followed immediately by washing the membranes with the reaction buffer (50 mM NaCl, 25 mM sodium phosphate (pH 7.5), and 5 mM BME).

After spotting, the membranes were air-dried and subjected to Typhoon visualization to detect Cy5 fluorescence at 633 nm. Quantitative analysis of the blot intensities for each membrane was performed using ImageLab software. The dissociation constants (K_d_) for the RNA-protein interactions were calculated using the Hill equation 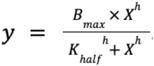 in Prism software, based on two or three replicates collected for each binding assay ^69^.

## 3. Results

### 3.1 RBM15 binds stem-loop structured RNA motifs

We focused our attention on characterizing the structural mechanism by which RBM15 interacts with RNA. For that, we focused our attention on characterizing interactions between RBM15 and lncRNAs for which secondary (2D) structures, supported by experimental restraints, are available.

We mined selective 2’ hydroxyl acylation analyzed by primer extension (SHAPE) and dimethyl sulfate (DMS) mutational profiling (MaP) chemical probing data available from RNA Atlas of Structure Probing (RASP) to fold the structures of several lncRNAs that have been shown to bind RBM15 in crosslinking and immunoprecipitation (CLIP) experiments (**Tables S1, S2**) ^6,31,32^. These lncRNAs include the X-Inactive Specific Transcript A-repeats (XIST A-repeats), the Functional Intergenic Repeating RNA Element Repeating RNA Domain (FIRRE RRD), Just Proximal to XIST (JPX), Five Prime to XIST (FTX), Long Intergenic Non-Protein Coding RNA 00630 (LINC00630), MicroRNA 503 Host Gene (MIR503HG), Long Intergenic Non-Protein Coding RNA 00894 (LINC00894), Inactivation Escape 1 (INE1), and Zinc Finger 674 Antisense RNA 1 (ZNF674-AS1). Given that many of these lncRNAs possess several isoforms, we focused on the splice variants that had the most chemical probing data coverage (number of nucleotides showing reactivity versus total number of nucleotides). Coverage varied from 8% to 100% across the lncRNAs. Details regarding the coverage of chemical probing data used to fold the structures are listed in **Table S2**.

The secondary structure predictions reflect intricately folded states for each of the nine lncRNAs, both in the *in vitro* and in-cell datasets (**Figures S2-10**). Overall, the chemical probing data is in good agreement with the reported structures, though it is noteworthy that the top three structures predicted for each RNA differ by less than 1 kcal/mol in free energy, suggesting local structural dynamics ^33–35^. However, insufficient coverage limited the application of deconvolution algorithms to effectively analyze the structural ensemble. Both *in vitro* and in-cell chemical probing data were available for XIST, FTX, and JPX; a comparison of the 2D structures from the two conditions reveals that each RNA folds differently. Structural differences are also observed in cases where the lncRNA is localized in the nucleus, cytoplasm, or chromatin. This is likely due to the highly crowded cellular environment with macromolecules and RNA binding events occurring in the cell ^36,37^.

Considering that our focus was on understanding how RBM15 interacts with RNA, we turned our focus to regions of the structured lncRNAs that interact with RBM15 as identified through CLIP experiments. Motif analysis revealed these regions typically possess CU, GA, and CA-containing sequences in stem-loop structured elements, accounting for approximately 85% of the identified interactions (**Tables S3, S4, Figure S11)**, regardless of their localization in the cell (**Figure 2A, 2B, Table S1**).

**Figure 2.**
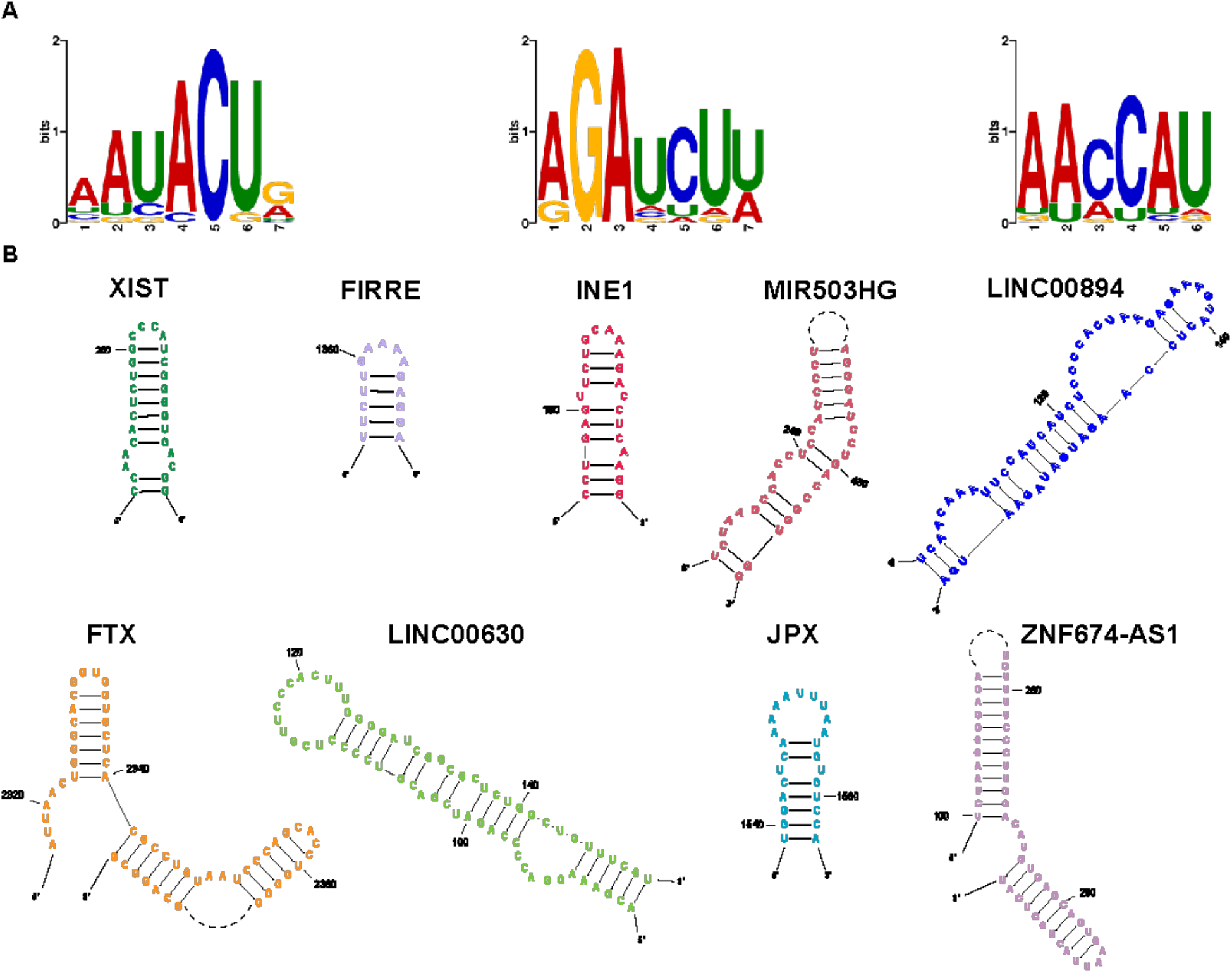
Stem-loop structured domains bound by RBM15. A) Binding sequence motif logo for RBM15 based on CLIP data. The logos highlight the predominant sequence motifs, including CU, GA, and CA, that are commonly found in stem-loop structures. B) RBM15 binding motifs were identified using CLIP data, supported by RNA 2D chemical probing experiments. A representative stem-loop (see also **Figure S11**) is shown for each lncRNA.

### 3.2 RBM15 RRM domains 2 and 3 facilitate binding with RNA

Using catRAPID, a computational tool used to predict RNA-protein interactions, we investigated which region of RBM15 is predicted to interact with RNA ^38^. As expected, the RRMs (RRM1: 170-252, RRM2: 374-451, and RRM3: 455-529), but not the SPOC domain, are predicted to interact with RNA (**Figure S12**). We next assessed the interaction between the RBM15 RRM domains and the lncRNAs. The catRAPID Graphic tool was used for RNAs shorter than 1,200nts, including the XIST A-repeats, the FIRRE RRD, INE1, MIR503HG, LINC00894, and FTX. For RNAs longer than 1,200nts, including LINC00630, JPX, and ZNF674-AS1, the catRAPID Fragments tool was employed ^39,40^. In agreement with the characteristics of typical RNA recognition motifs, regions corresponding to the RRM domains of RBM15 are predicted to bind with the lncRNAs (**Figure 3**). Interestingly, we observed that RRMs 2 and 3 are predicted to bind preferably over RRM1 for the majority of lncRNAs as indicated by greater distribution of red in the heat maps (**Figure 3**).

**Figure 3.**
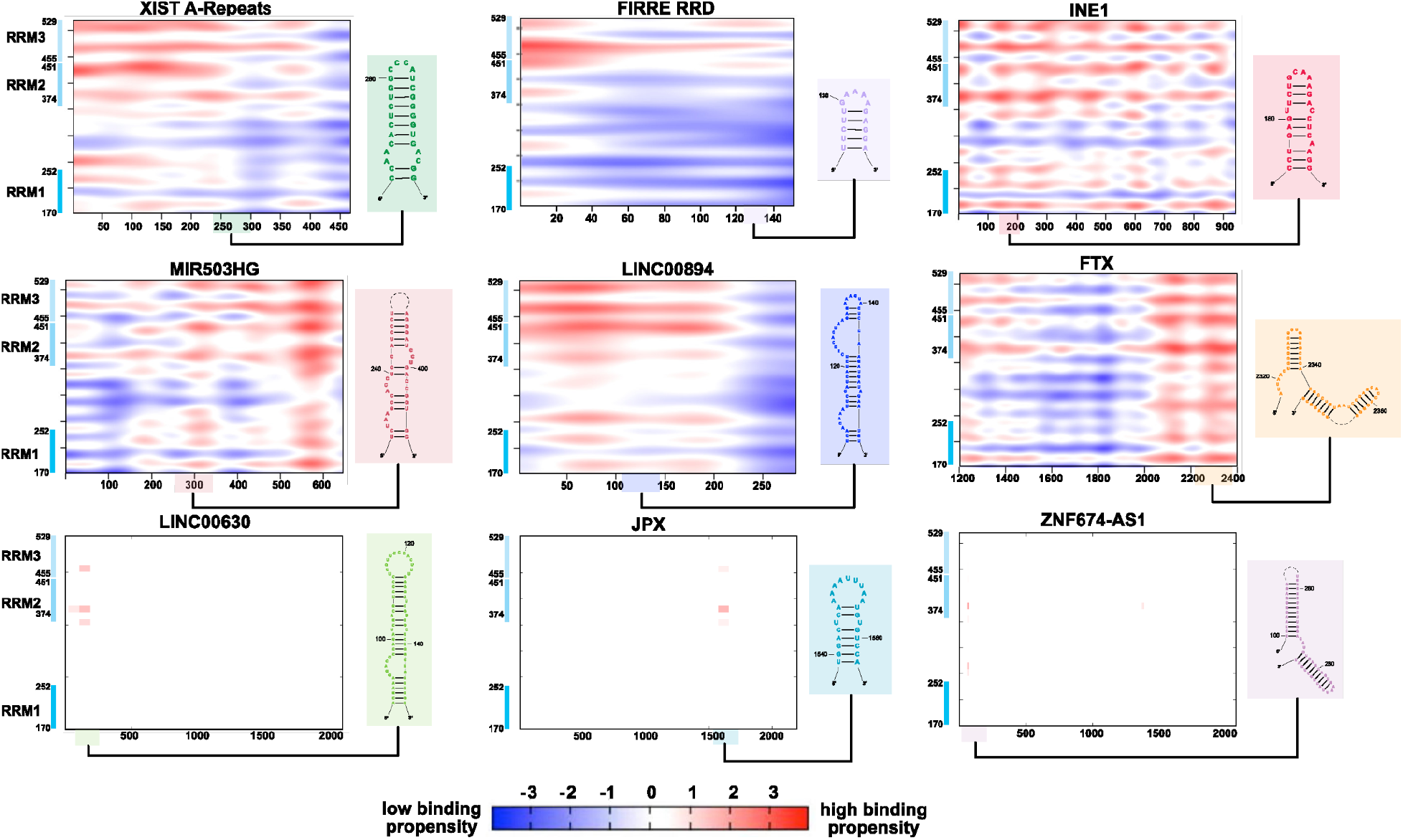
catRAPID predicted interaction domains for RBM15 with lncRNAs. catRAPID predictions of RBM15 interaction domains with lncRNAs are shown. Interaction propensity is indicated by color: blue for weak interactions, and red for high interactions. The RRM domains of RBM15 are indicated with blue vertical lines along the y-axis. Nucleotides and select 2D structured elements corresponding to RBM15 binding regions, are highlighted on the x-axis.

Notably, the nucleotides predicted by catRAPID to bind to RBM15 are moderately analogous to the RBM15 CLIP sites, encompassing regions possessing stem-loop structured motifs (**Figure 3**). For example, the lncRNAs MIR503HG and FTX are predicted to interact with RBM15 in the middle and at the 3’ end of the RNA, respectively. Investigation of the 2D structures in these regions corresponds to the stem-loop structured motifs identified in our CLIP-2D structure analysis. The locations of some of the stem-loops identified by CLIP and catRAPID are not in as strong agreement, such as with the XIST A-repeats. This may be due to false positives in the CLIP experiments or limitations associated with catRAPID. Specifically, the catRAPID algorithms employed are trained on RNA sequence and structural data, but the secondary structure information regarding the RNA is determined *in silico* through RNAfold in the Vienna package as opposed to chemical probing-supported structures ^38^. In the absence of experimental structure constraints, structures predicted in catRAPID may not coincide with experimentally determined structures. RBM15 may bind and recognize both sequence and structured elements, and in the absence of experimentally validated structural training data, these may have been mispredicted. Nevertheless, the overall consistency between the two methods confirms the interaction of RBM15 with each of these lncRNAs.

Of the nine lncRNAs in this study, the two most functionally characterized are the lncRNAs XIST and FIRRE. Thus, we used these two RNAs to experimentally validate the catRAPID and CLIP results. Considering the CLIP and chemical probing data supported the notion that stem-loop structured RNAs are bound by RBM15, we centered our investigation on the stem-loops from XIST A-repeats and FIRRE RRD lncRNAs (see also **Figures 2 and 3**) and their interaction with individual (RRM1, RRM2, and RRM3) and tandem (RRM12, RRM23, RRM123) RRM domains from RBM15. The 32nt stem-loop from the XIST A-repeats consists of a 2nt bulge followed by 8bps leading to a 4nt apical loop. On the other hand, the 19nt stem-loop from the FIRRE RRD consists of a 5bp stem leading to a 5nt apical loop. Neither stem-loops consist of the DRACH motif, however, both are adjacent to single-stranded DRACH motifs: GAACC and AAACU for XIST and FIRRE stem-loops respectively.

To dissect the individual and combined contributions of each RNA recognition motif (RRM) within our protein of interest, we performed RNA filter dot blot assays using various constructs comprising different combinations of RRM domains ^41^. Individually, RRM1 bound the target RNAs with moderate affinities (K_d_ = 433 and 370 nM for the XIST and FIRRE stem-loops, respectively), while RRM2 exhibited substantially weaker interactions (K_d_ > 1.5 µM for both RNAs). In contrast, RRM3 showed the highest individual affinity for RNA, with a K_d_ of ~100 nM for both RNAs (**Figure 4A, Figure S13A**).

**Figure 4.**
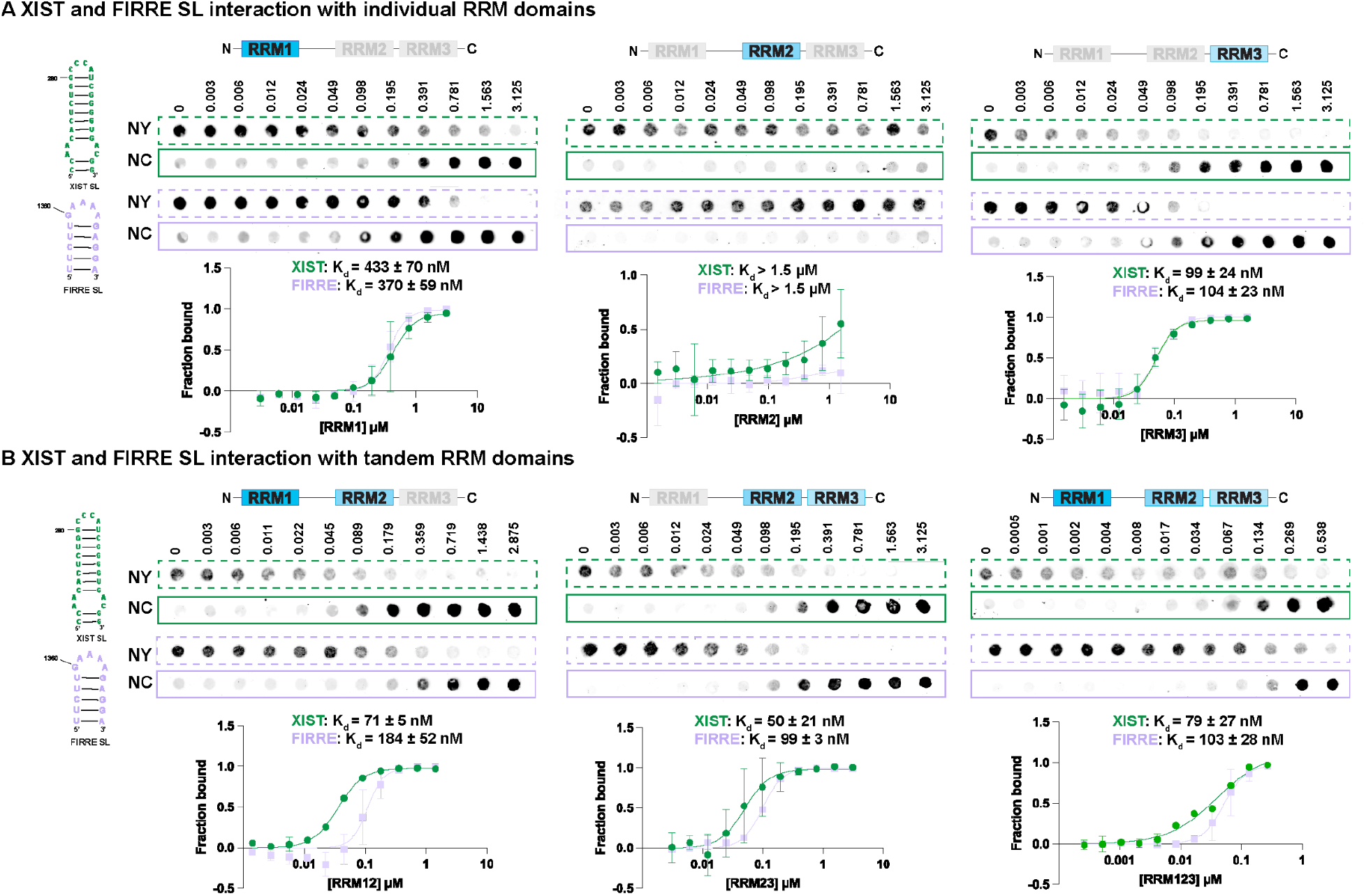
RBM15 interaction with the CLIP-SHAPE stem-loops. A) Dot blot interaction analysis of the XIST A-repeats stem-loop (32nt; green) and the FIRRE RRD stem-loop (19nt; purple) with individual and B) tandem RRM domains. NY = nylon, NC = nitrocellulose, SL = stem-loop. Binding affinities and error bars from replicates are presented in the fitted plots. Protein concentrations are indicated on top (µM).

We next assessed RNA binding by tandem RRM constructs to examine whether domain combinations enhance affinity (**Figure 4B, Figure S13B**). A construct encompassing RRM1 and RRM2 along with the intervening 123-amino acid linker (RRM12) bound the XIST and FIRRE RNAs with improved affinities (K_d_ = 71 nM and 184 nM, respectively), suggesting synergistic interactions between RRM1 and RRM2 or a potential scaffolding effect of the long linker that optimally positions RRM1 for RNA engagement. Notably amino acids comprising this linker include several residues known to facilitate interactions with RNA through hydrogen bond and pi-pi stacking interactions with RNA. These comprise of 8 aromatic residues (2 phenylalanines and 6 tyrosines), and 40 hydrogen bonding amino acids including arginines, lysines, glutamic/aspartic acids, asparagines and glutamines. Despite RRM2’s weak affinity alone, its inclusion with RRM1 clearly enhances the overall binding, indicative of functional cooperativity.

A separate construct containing RRM2, the short 3-amino acid linker, and RRM3 (RRM23) exhibited even stronger binding to XIST (K_d_ = 50 nM) and FIRRE (K_d_ = 99 nM), likely due to the coaxial stacking of RRM2 and RRM3 predicted by AlphaFold (**Figure 5, Figure S14**), which may promote an extended RNA-binding interface or stabilize a high-affinity conformation. Notably, despite RRM2’s poor binding individually, its structural integration with RRM3 results in significantly enhanced RNA affinity.

**Figure 5.**
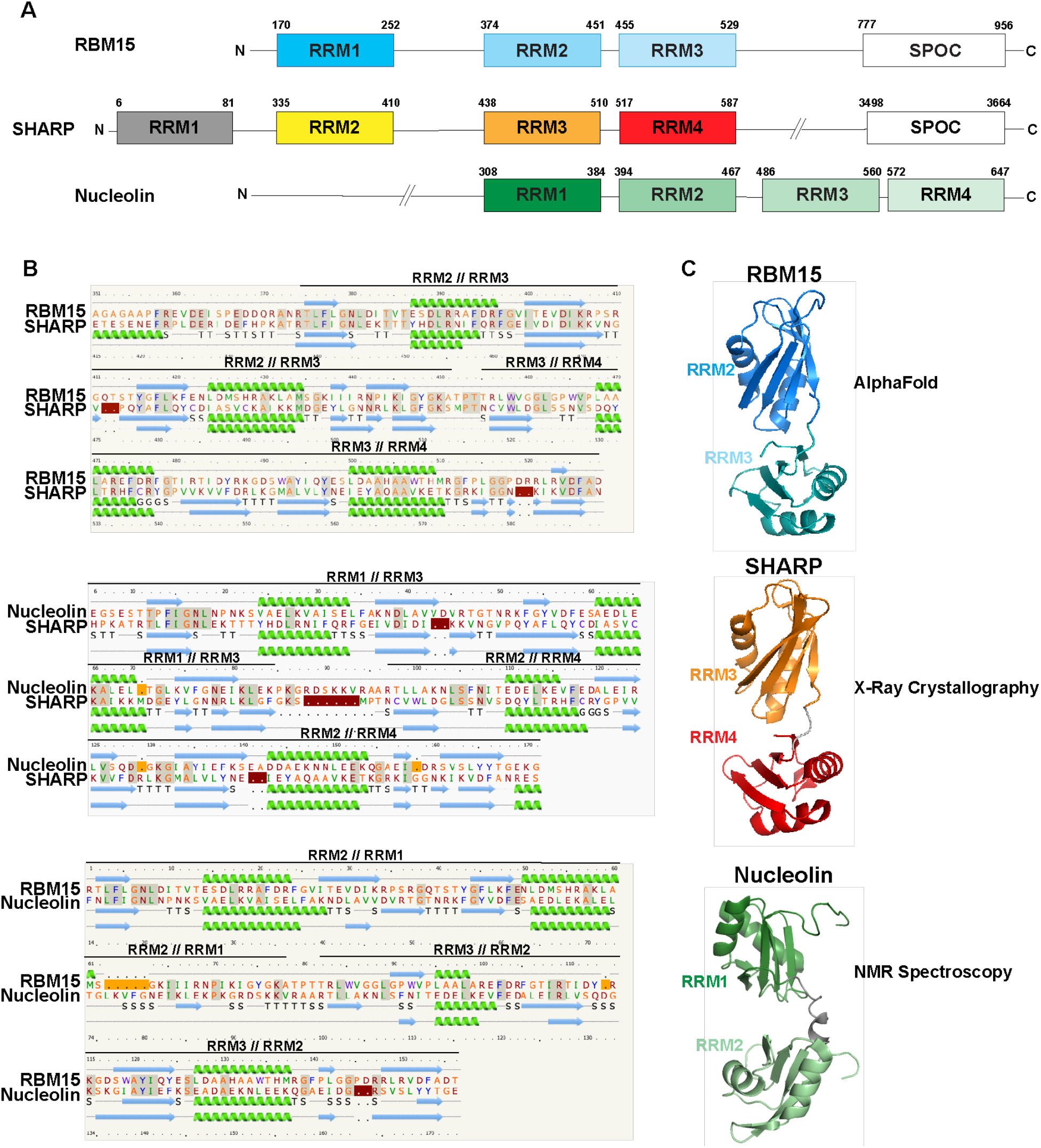
Structural comparison of RBM15, SHARP, and Nucleolin RRM domains. A) Aligned RBM15, SHARP, and Nucleolin protein sausage plots. B) Sequence and secondary structure comparison of RBM15 RRMs 2 and 3, SHARP RRMs 3 and 4, and Nucleolin RRMs 1 and 2, aligned using Phyre2. Red/orange patches indicate amino acids present/absent between the compared protein sequences. C) 3D structural comparison between RBM15 RRM23 (AlphaFold), SHARP RRM34 (PDB ID 4P6Q), and Nucleolin RRM12 (PDB ID: 1FJE) tandem constructs.

The full-length construct containing all three RRMs (RRM123) bound the XIST RNA with a K_d_ of 79 nM (FIRRE was comparable with a K_d_ of ~100 nM), values that are comparable to RRM12 but slightly weaker than RRM23. This suggests that while all three RRMs contribute to RNA recognition, the cooperative advantage between RRM2 and RRM3 may be partially offset when RRM1 is included, possibly due to increased conformational flexibility or suboptimal spatial arrangement imposed by the long linker between RRM1 and RRM2.

Together, these results demonstrate that the RNA-binding affinity of the protein is not solely determined by individual RRM affinities but is significantly influenced by domain arrangement, inter-domain linkers, and structural cooperativity.

### 3.3 RBM15 RRM23 Crafts a ‘Sandwich-Like’ Structure to Bind RNA

Our experimental analysis revealed that although RBM15 RRM2 does not efficiently bind RNA while RRM3 does, a tandem RRM23 construct improves association with RNA. We hypothesized that the two RRMs may interact, adopting a structural orientation that is required to form a stably structured RNA complex. Interestingly, RRMs 2 and 3 are separated by just three amino acids, an indication that they may share a structural interface.

We used Phyre2 to predict and analyze the structure of the RBM15 protein based on homology modeling to gain more insight into how it is organized in solution ^42^. Phyre2 identified a known homolog of RBM15, SMRT/HDAC1 Associated Repressor Protein (SHARP), and Nucleolin; these proteins each possess four RRMs ^43,44^. RBM15 RRMs 1, 2, and 3 align with SHARP RRMs 2, 3, and 4; Nucleolin RRMs 1 and 2 align with RBM15 RRMs 2 and 3, respectively. (**Figure 5A**).

Alignments of the sequences reveal good agreement (**Figure 5B**). Furthermore, the Phyre2 transitive consistency score for multiple sequence alignment reveals excellent scores (out of 100) for each conserved RRM set: RBM15 RRM1-SHARP RRM2, RBM15 RRM2-SHARP RRM3, RBM15 RRM3-SHARP RRM4 have scores of 80, 94, and 98, respectively. RBM15 RRMs 2 and 3 scored 96 and 88 relative to Nucleolin RRMs 1 and 2, while SHARP RRMs 3 and 4 scored 75 and 84 compared to Nucleolin RRMs 1 and 2, indicating strong alignments among the three proteins ^45^.

Interestingly, the crystal structure of SHARP RRM234 reveals that RRMs 3 and 4 form a heterodimer, with these domains separated by just seven amino acids (PDB ID: 4P6Q) ^43^. Similarly, the NMR structure of Nucleolin RRM12 reveals that RRMs 1 and 2 interact, with these domains separated by only ten amino acids (PDB ID: 1FJE) ^44^. Inspired by this, we used AlphaFold3 to model the 3D structure of RBM15. This revealed a model whereby RRMs 2 and 3 are coaxially stacked, and RRM1 is folded independently of the rest of the structure (**Figure 5C, Figure S14**) ^46^. This is similar to the experimentally determined 3D structures of SHARP and Nucleolin RRM domains, indicating a rationale and mechanism for RBM15’s recognition and binding ^43,47^.

## 4. Discussion

While several genome-wide crosslinking studies have been carried out to identify sites of protein binding with RNA, knowledge of the molecular mechanisms by which these interactions occur is lacking. In this study, we mined chemical probing data available from the RASP database to investigate the structure of several lncRNAs and identified RBM15 sites of binding throughout these transcripts based on CLIP data from the ENCODE and POSTAR3 databases. We then computationally and experimentally validated these findings using stem-loops from two lncRNAs, the XIST A-repeats, and the FIRRE RRD.

We find that RBM15 binding sites occur in stem-loop structured regions across each of the lncRNAs investigated. These regions typically include CU, GA, and CA-containing sequences. This is different from other reported RBM15 binding motifs, which suggest RBM15 interacts with U-rich RNAs ^48^. Here, these studies focus primarily on sequence motifs, whereas we focus on sequences encompassed in structured domains. RBM15’s RRM3 appears to be important in binding the RNAs investigated in this study, as it is the strongest binder among individual RRMs and also improves the binding affinity when in a tandem RRM construct. Our results also reveal that RRMs 2 and 3 form a heterodimer and are primarily responsible for facilitating RBM15’s interaction with RNA. This is similar to what is observed for its homolog, SHARP, and another RBP, Nucleolin, which binds RNA through two of its coaxially stacked RRMs ^43,44^. Using *in silico* approaches, we find that RBM15 interacts with RNA stem-loops through a mechanism akin to that of Nucleolin: RRM23 forms a defined pocket that serves as a stable binding site for the RNA, with the RRM2 and RRM3 motifs working in concert to ensure increased affinity.

The functional role of the interactions between RBM15 and the majority of these lncRNAs has yet to be extensively explored; however, RBM15 is known to play a key role in the recruitment of m6A writer complex to RNA to facilitate the N6 methylation of adenosine ^6,49^. Though these adenosines are nested in DRACH sequences, not all adenosines nested within DRACH sequences are modified. RBM15, which has specific binding properties with RNA, is believed to regulate which DRACH sequences harbor adenosines that will be modified ^50^. Our results suggest that this is accomplished through its affinity for CU, GA, and CA-containing sequences in stem-loop motifs that are either proximally located or structurally adjacent to DRACH sequences and m6A modifications (**Figure 6, Figure S15**). While further investigation is needed to fully comprehend how RBM15 is implicated in the regulatory roles of the lncRNAs that we focused on in this study, our results nevertheless provide structural insight into how RBM15 recognizes and binds structured regions of RNAs.

**Figure 6.**
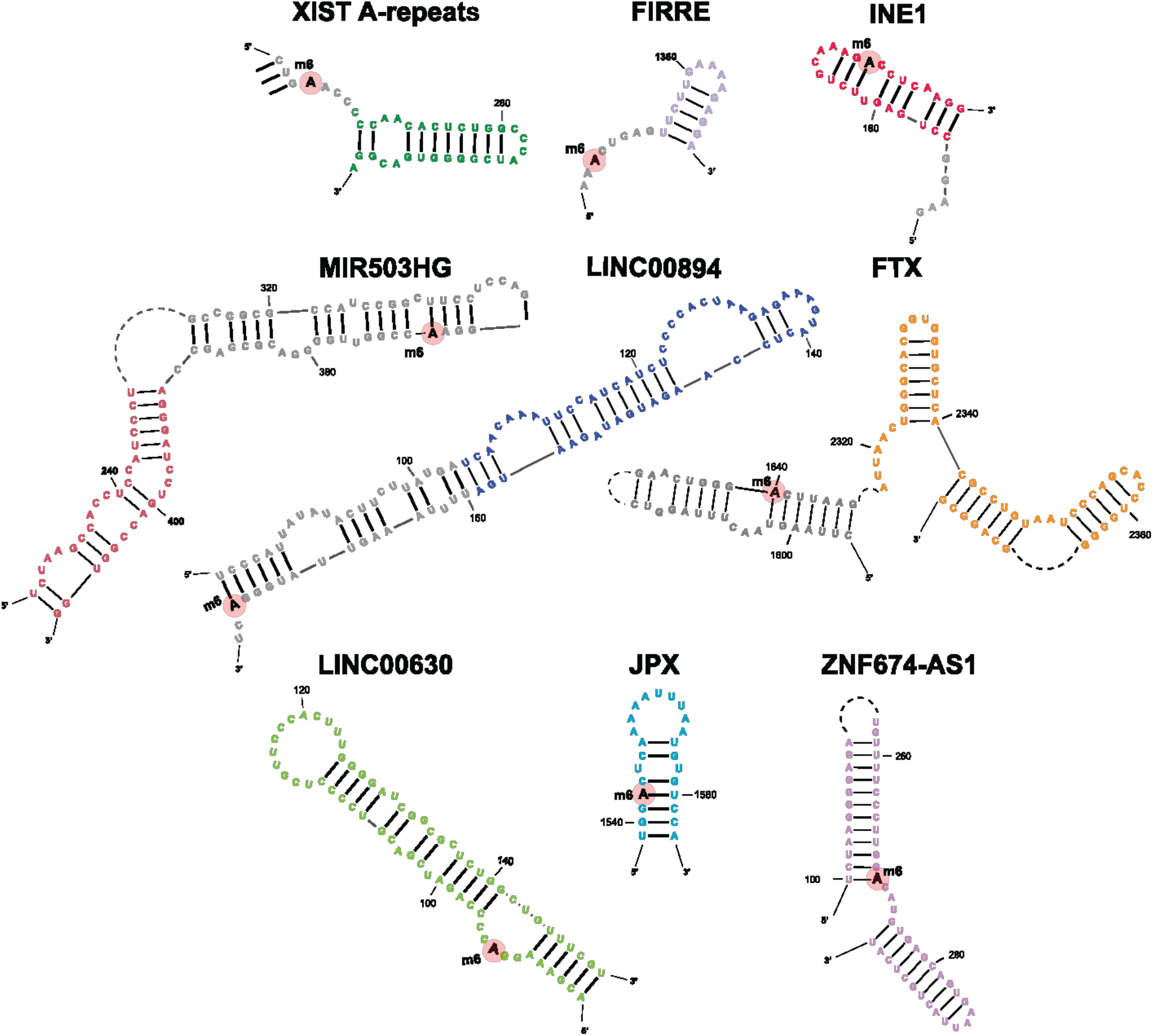
lncRNA m6A enrichment sites in RBM15 binding stem-loops. DRACH-m6A enrichment sites on RBM15 CLIP-SHAPE hairpin structures. m6A sites are highlighted with pink circles, and RBM15 CLIP sequences are colored; grey nucleotides correspond to regions outside of the CLIP data. See also Figure 2 and Figure S15.

## Data Availability

All data is included in this manuscript.

## Acknowledgments

The authors would like to thank members of the Jones lab for useful discussions and the Sattler lab for RBM15 plasmids.

## Author Contributions

EB, CM, SX, YPOS, MS, LB, FL, JJ, TI, DM, DK, JW, VZ, LV, JVH, GA, KB, LZ, and ANJ carried out data mining, folding of RNA structures, and bioinformatic analysis. ANJ, EB, LE, CB, BE and ST prepared samples and carried out interaction assays and analyses. GW wrote scripts for assessing data. EB and ANJ wrote the manuscript and EB, BE, and ANJ revised the manuscript. All authors reviewed the manuscript.

## Funding and Additional Information

This work was funded by the National Science Foundation, 2243667 to A.N.J.

