## SupplementalData for "Bioinformatic and Experimental Characterization of the RBM15 RNA Binding Protein"

**Supplementary Figures**

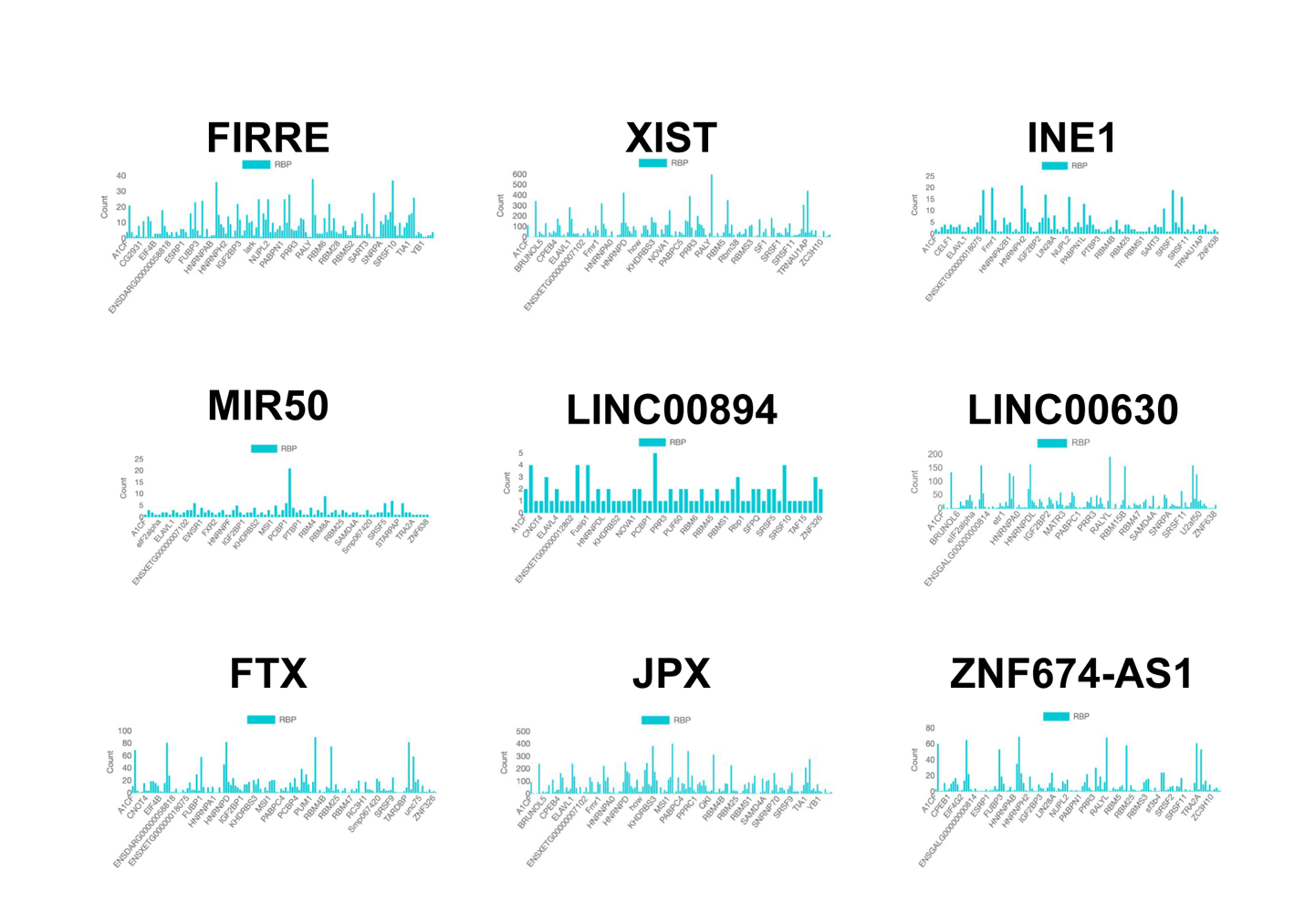

**Figure S1.** oRNAment bar plots for RBPs predicted to interact with the lncRNAs.

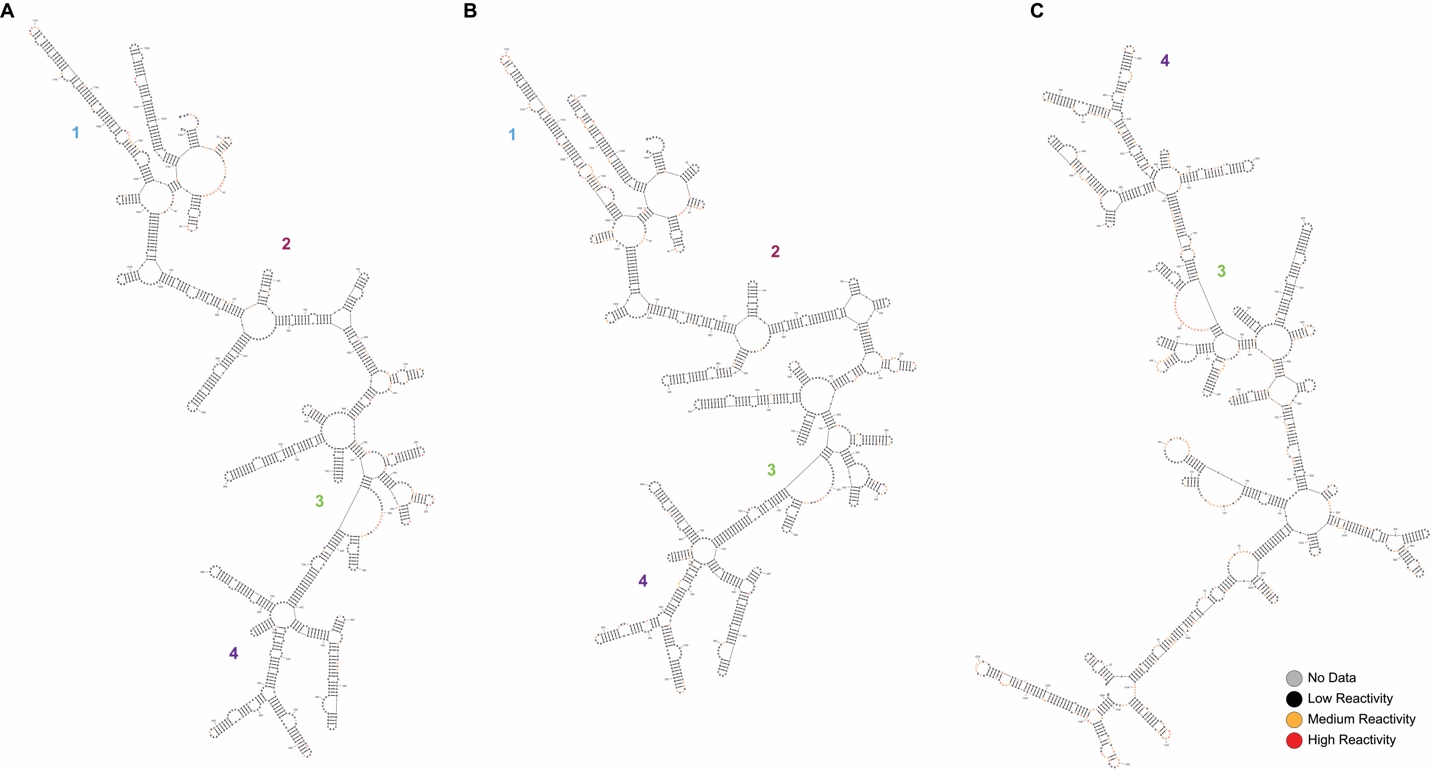

**Figure S2.** Secondary structures of the lncRNA X-Inactive Specific Transcript A-repeat (XIST A-repeats) (*in vivo*) based on NAI-N3 SHAPE chemical probing data A) in the cytoplasm B) in the nucleoplasm, and C) in chromatin. Numbers correspond to structured regions that are maintained despite the localization of the transcript. Chemical probing data is mapped to each structure (gray = no data, black = low reactivity, orange = medium reactivity, and red = high reactivity).

**Figure S3**. Secondary structure of the lncRNA Functional Intergenic Repeating RNA Element (FIRRE) (*in vivo*) based on DMS-MaPseq chemical probing data. The purple highlight indicates the single-stranded RNA template used in binding assays. Chemical probing data is mapped to each structure (gray = no data, black = low reactivity, orange = medium reactivity, and red = high reactivity).

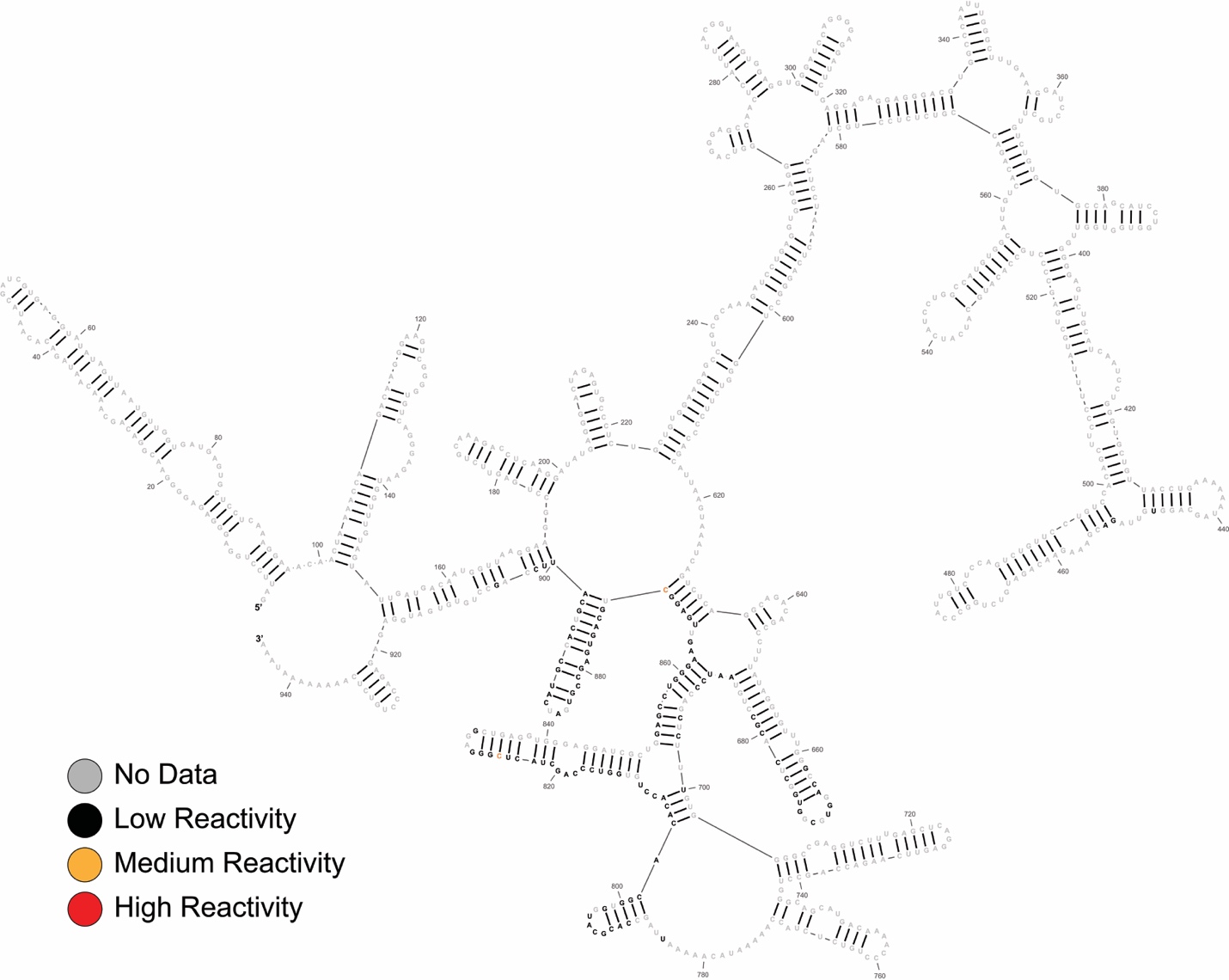

**Figure S4.** Secondary structure of the lncRNA Inactivation Escape 1 (INE1) based on *in vivo* DMS-MaPseq chemical probing data. Chemical probing data is mapped to each structure (gray = no data, black = low reactivity, orange = medium reactivity, and red = high reactivity).

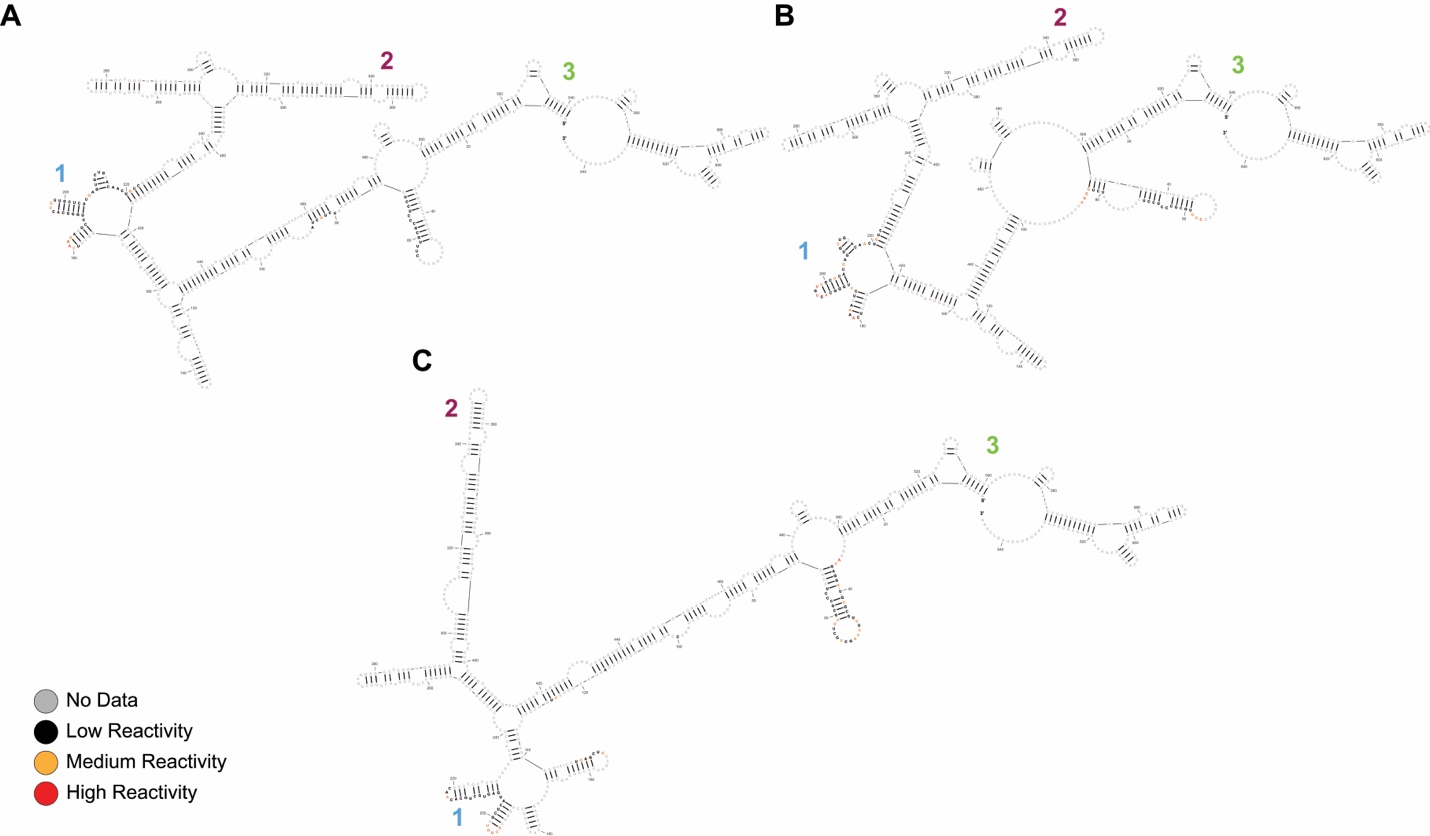

**Figure S5.** Secondary structure of the 5-way multiway junction supported by NAI-N3 SHAPE chemical probing data for the lncRNA MicroRNA 503 Host Gene (MIR503HG). The 5-way junction A) in chromatin *in vitro* and B) in chromatin *in vivo* are structurally identical. The 5-way junction predicted for MIR503HG in the nucleoplasm C) only shares one helical domain (3) with structures derived from MIR503HG when localized in chromatin, but similar structural elements are also observed between each condition (numbered 1 and 2). The. Chemical probing data is mapped to each structure (gray = no data, black = low reactivity, orange = medium reactivity, and red = high reactivity).

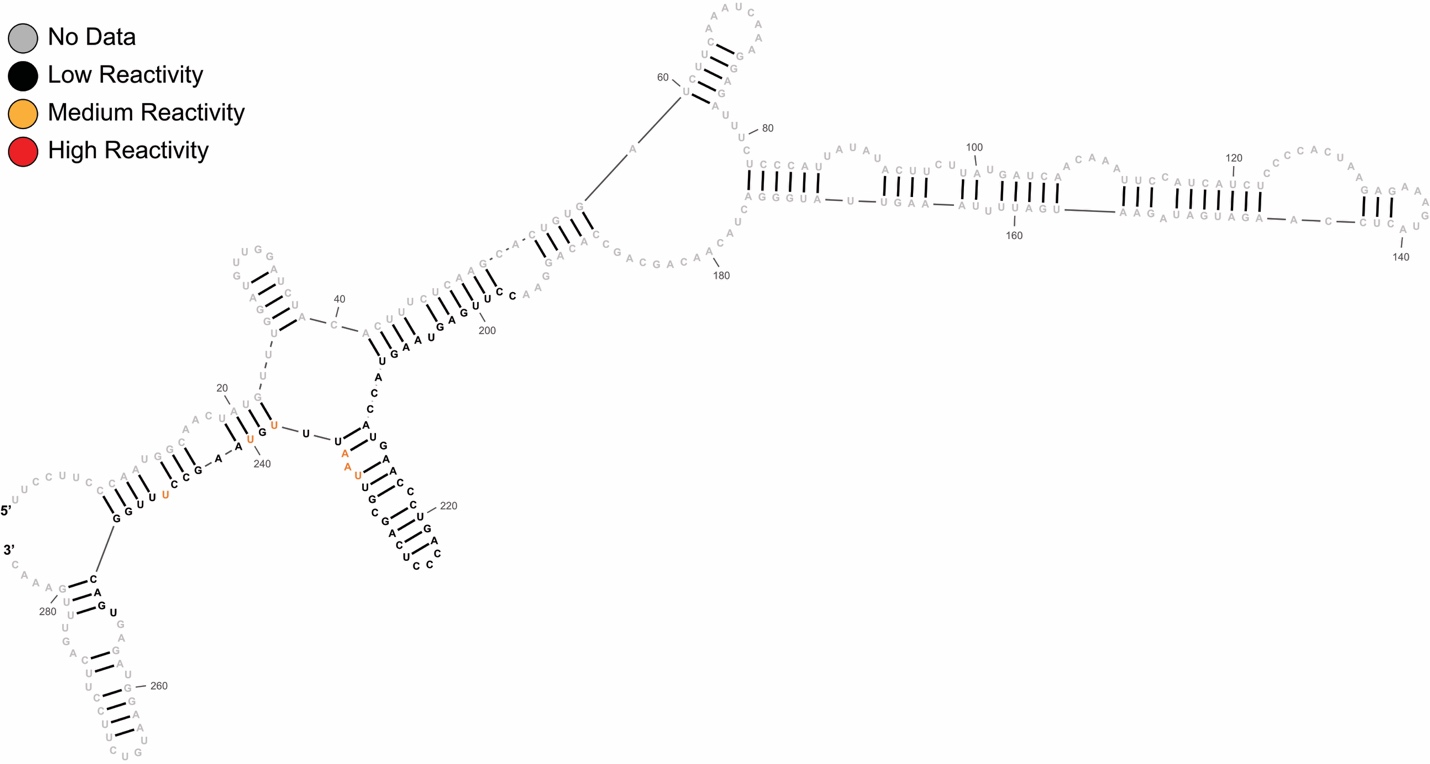

**Figure S6**. Secondary Structure of the lncRNA Long Intergenic Non-Protein Coding RNA 00894 (LINC00894) (*in vitro*) based on NAI-N3 SHAPE chemical probing data. Chemical probing data is mapped to each structure (gray = no data, black = low reactivity, orange = medium reactivity, and red = high reactivity).

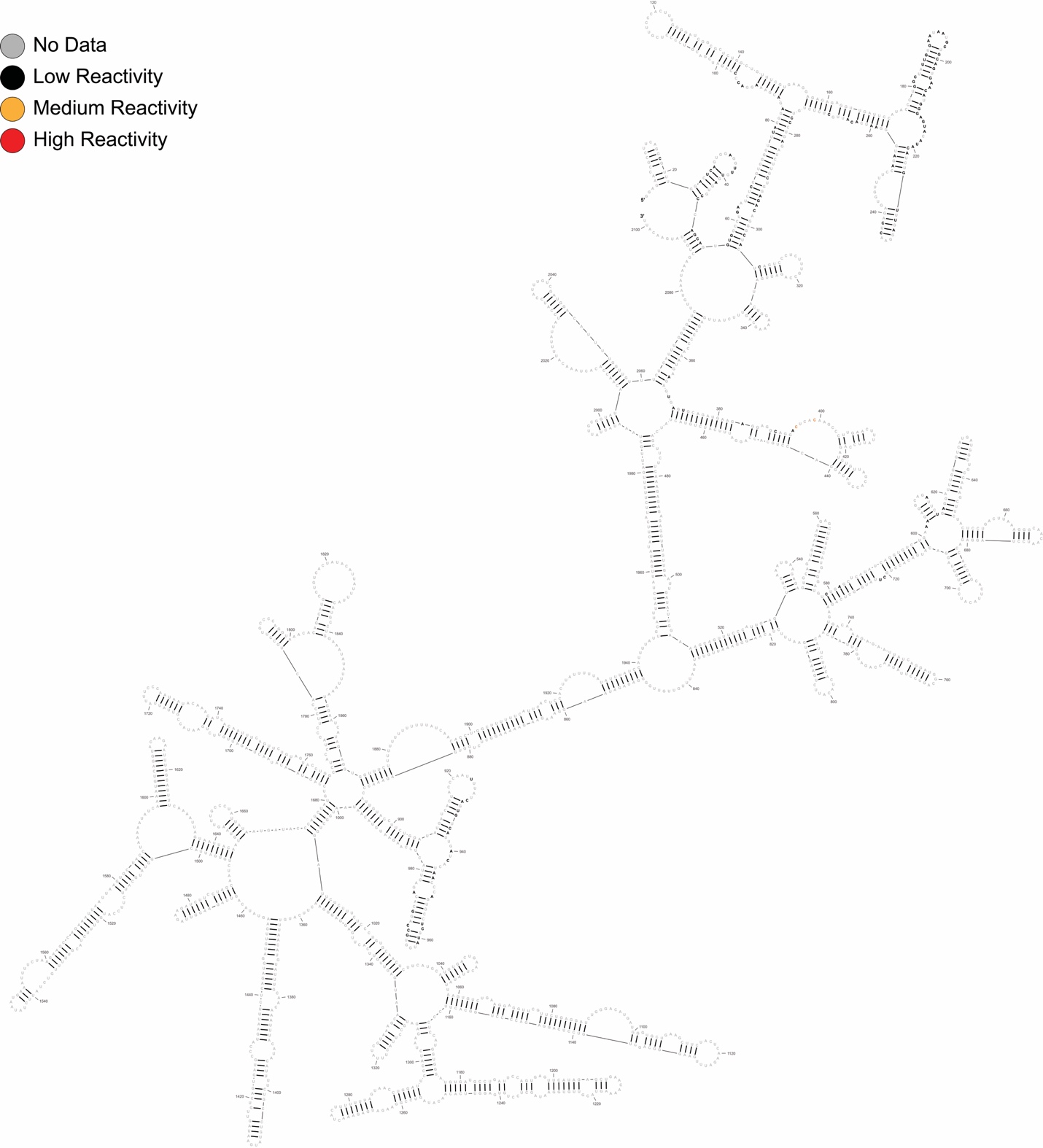

**Figure S7**. Secondary structure of Long Intergenic Non-Protein Coding RNA 00630 (LINC00630) (*in vivo*) based on DMS-MaPseq chemical probing data. Chemical probing data is mapped to each structure (gray = no data, black = low reactivity, orange = medium reactivity, and red = high reactivity).

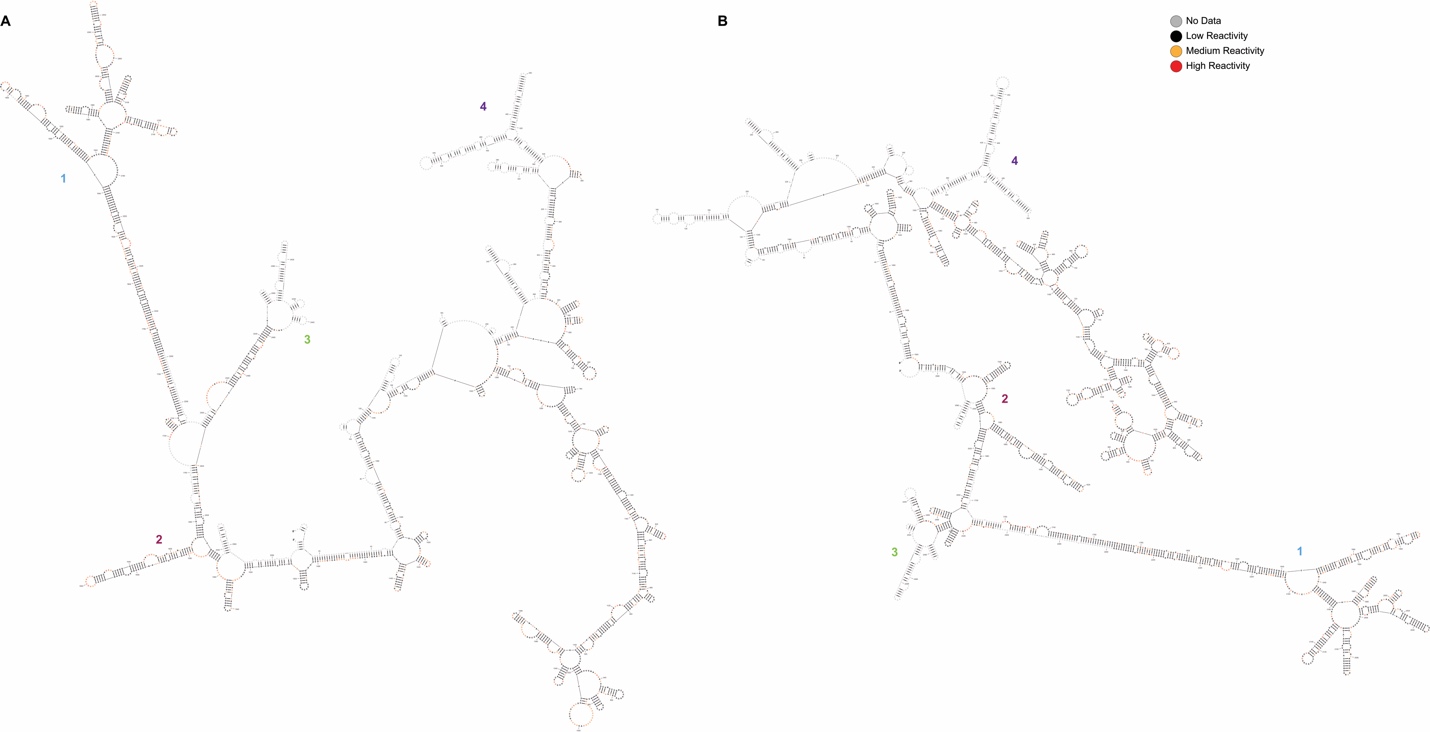

**Figure S8.** Secondary structure of the lncRNA Five Prime to XIST (FTX) based on NAI-N3 SHAPE chemical probing data. Four domains (1-4) are similar between the A) FTX chromatin *in vitro* and B) FTX chromatin *in vivo* predicted structures. Chemical probing data is mapped to each structure (gray = no data, black = low reactivity, orange = medium reactivity, and red = high reactivity).

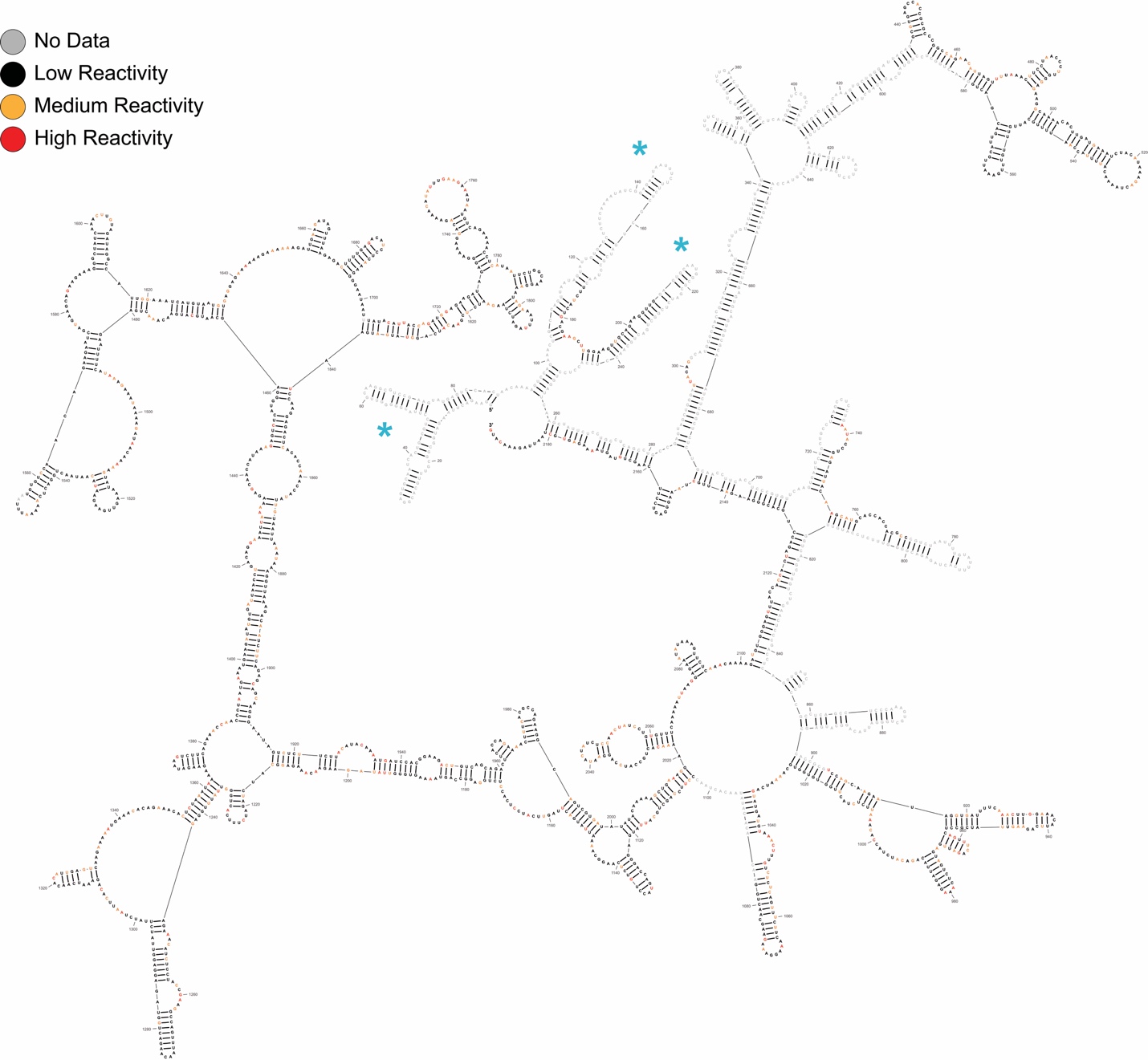

**Figure S9.** Secondary structure of the lncRNA Just Proximal to XIST (JPX) (chromatin *in vitro*) based on NAI-N3 SHAPE chemical probing data. Blue asterisks indicate structured domains with a similar fold to a previously reported structure corresponding to nucleotides 1 - 343 of JPX ^1^. Chemical probing data is mapped to each structure (gray = no data, black = low reactivity, orange = medium reactivity, and red = high reactivity).

**Figure S10.** Secondary structure of the lncRNA Zinc Finger 674 Antisense RNA 1 (ZNF674-AS1) (*in vivo*) based on DMS-MaPseq chemical probing data. The purple highlight indicates the dsRNA template used in binding assays. Chemical probing data is mapped to each structure (gray = no data, black = low reactivity, orange = medium reactivity, and red = high reactivity).

**Figure S11.** iCLIP RBM15 crosslinking RNA binding sites with each lncRNA.

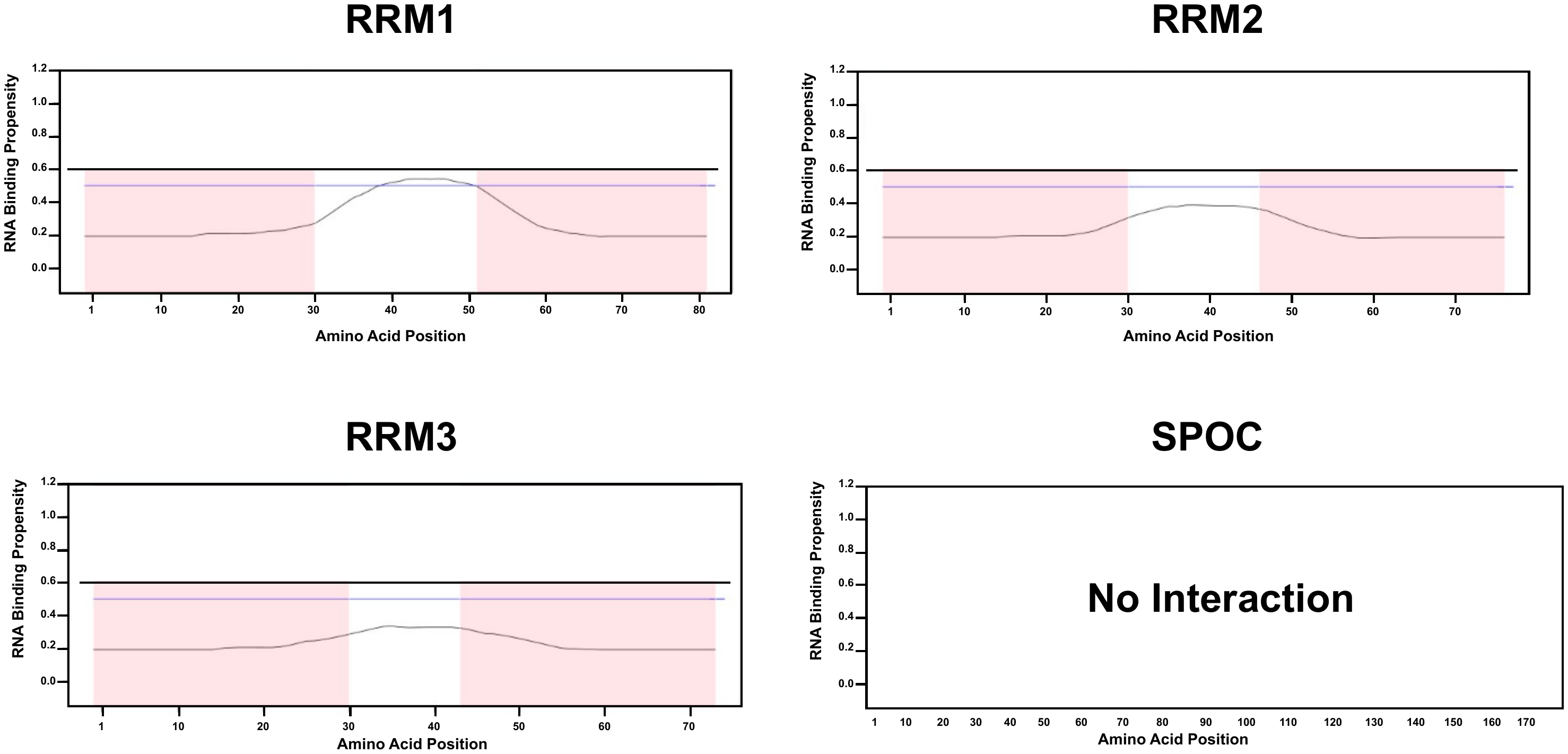

**Figure S12**. **catRAPID Interaction Domains for RBM15 with lncRNAs.** catRAPID predictions for the interaction of individual RBM15 domains with lncRNAs are illustrated. An RNA-binding propensity above 0.6 (horizontal line) is considered significant. The x-axis represents each amino acid, while the y-axis shows the predicted RNA-binding propensity.

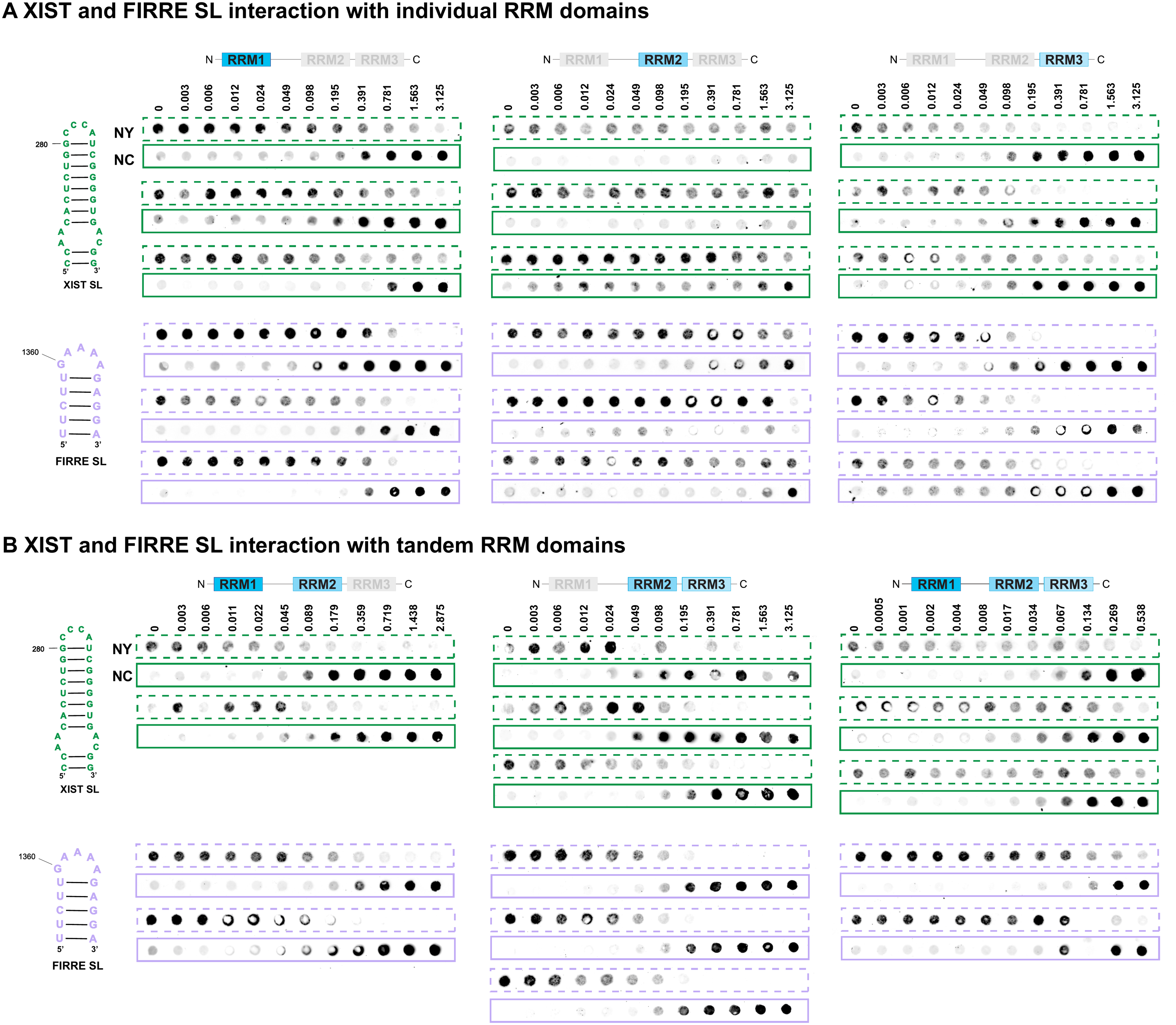

**Figure S13. RBM15’s interaction with the iCLIP stem-loops**. Quantitative replicate of dot blot interaction analysis of XIST A-repeat stem-loop (32nt; green) and FIRRE RRD stem-loop (19nt; purple) with A) individual RRM and B) tandem RRM domains. NY = nylon, NC = nitrocellulose, SL = stem-loop. Protein concentrations are indicated on top (µM).

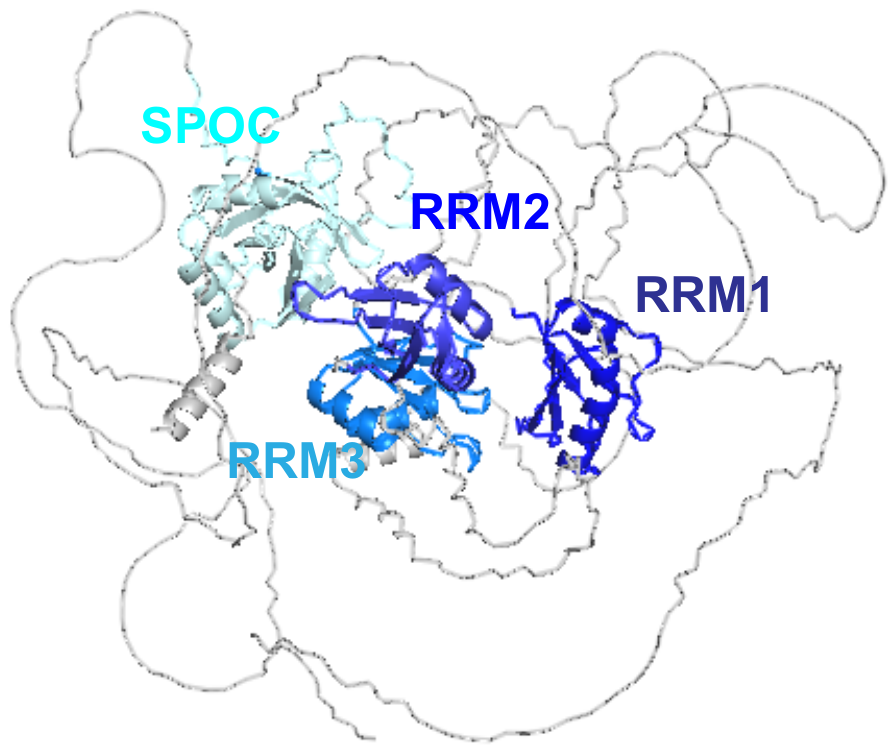

**Figure S14.** AlphaFold rendering of RBM15. RRMs and SPOC domains are labeled and shown in shades of blue.

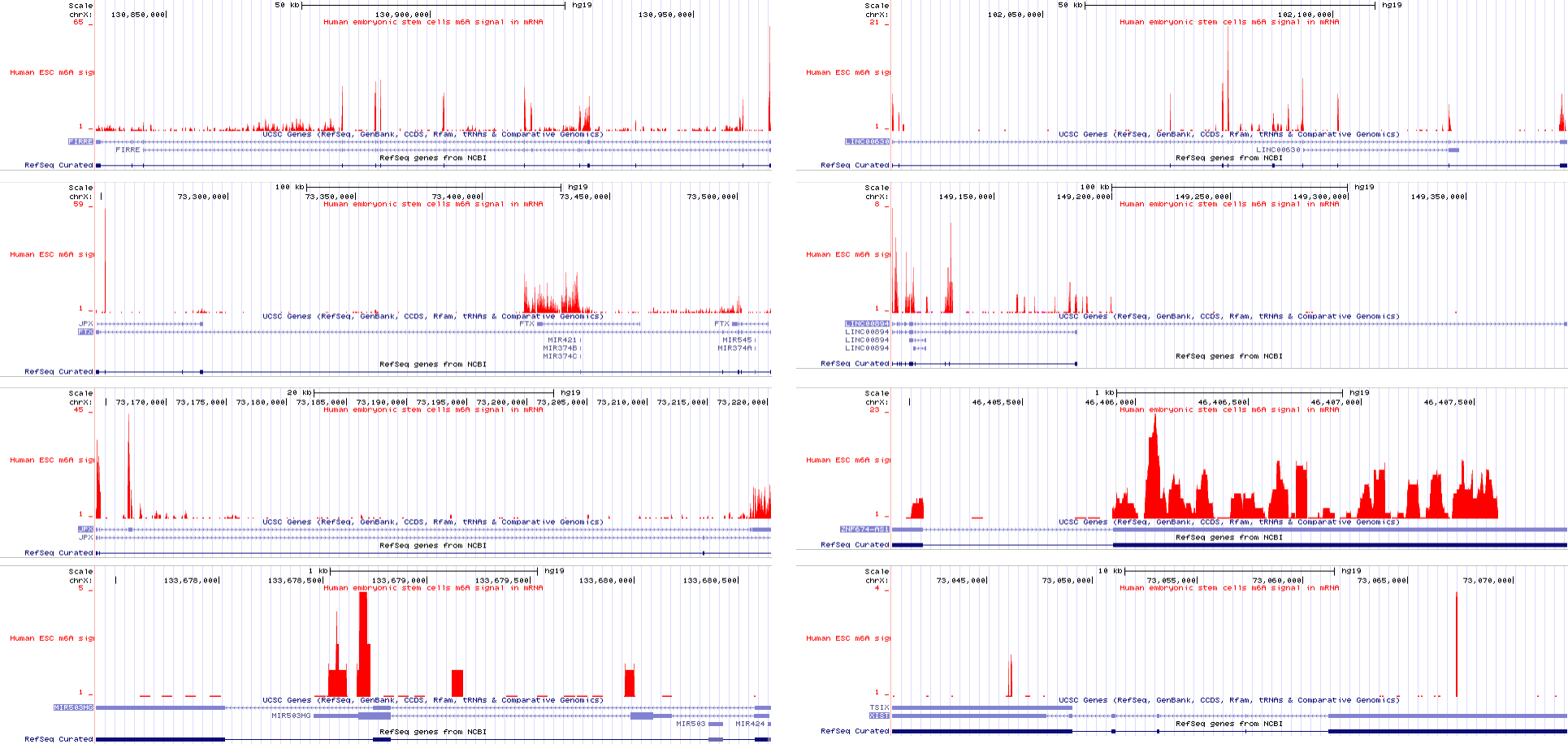

**Figure S15**. **m6A enrichment of the 9 lncRNAs investigated in this study**. The m6A enrichment data was sourced from the UCSC genome browser using the human embryonic stem cells m6A signal in the mRNA track setting.

**Supplementary Tables**

**Table S1.** CLIP RBM15 binding sites

| **RNA and ID** | **Chromosome Location** | **Technology and Peak Calling Method** | **Data Accession** |
| --- | --- | --- | --- |
| XIST (-)  ENST00000429829 | chrX:73851205-73851245;-  chrX:73851265-73851325;-  chrX:73851345-73851425;-  chrX:73851445-73851465;-  chrX:73851525-73851605;-  chrX:73851625-73851665;-  chrX:73851685-73851825;-  chrX:73851845-73852105;-  chrX:73852125-73852145;-  chrX:73852165-73852345;-  chrX:73852365-73852705;- | iCLIP: Piranha | GSE78030,GSM2064710 |
| FIRRE (-)  ENST00000427391 | chrX:131830443-131830444;-  chrX:131830367-131830368;-  chrX:131830312-131830332;-  chrX:131830307-131830360;-  chrX:131783901-131783916 | iCLIP: CITS, PureCLIP  eCLIP ENCODE | GSE78030,GSM2064710  ENCODE: ENCFF739LLZ, ENCFF086HIT |
| INE1 (+)  ENST00000456273 | chrX:47204936-47204969;+  chrX:47204997-47205020;+  chrX:47205091-47205110;+  chrX:47205203-47205234;+  chrX:47205215-47205252;+  chrX:47205314-47205353;+  chrX:47205514-47205535;+  chrX:47205653-47205691;+ | eCLIP | ENCODE: ENCFF739LLZ, ENCFF086HIT |
| MIR503HG (-)  ENST00000457876 | chrX:134545649-134545650;-  chrX:134544531-134544532;-  chrX:134544676-134544677;-  chrX:134544627-134544628;- | iCLIP: CITS, PureCLIP | GSE78030,GSM2064710 |
| LINC00894 (+)  ENST00000455777 | chrX:149960902-149960903;+  chrx:149962628-149962666;+ | iCLIP CITS  eCLIP | GSE78030,GSM2064710  ENCODE: ENCFF739LLZ, ENCFF086HIT |
| LINC00630 (+)  ENST00000440496 | chrX:102826990-102826991;+  chrX:102769217-102769247;+  chrX:102769268-102769300;+  chrX:102816997-102817022;+  chrX:102884250-102884287;+  chrX:102884951-102884971;+ | iCLIP: PureCLIP, Piranha, CITS  eCLIP | GSE78030,GSM2064710  ENCODE: ENCFF739LLZ, ENCFF086HIT |
| FTX (-)  ENST00000429124 | chrX:74280445-74280446;-  chrX:74281025-74281026;-  chrX:74278616-74278716;- | iCLIP: CITS, PureCLIP  eCLIP | GSE78030,GSM2064710  ENCODE: ENCFF739LLZ, ENCFF086HIT |
| JPX (+)  ENST00000414209 | chrX:73999356-73999357;+  chrX:73999532-73999533;+  chrX:73999257-73999258;+  chrX:74000101-74000102;+  chrX:74000146-74000147;+  chrX:74000339-74000340;+  chrX:73999220-73999221;+  chrX:73947342-73947343;+  chrX:73999046-73999047;+  chrX:73947331-73947332;+  chrX:73999848-73999849;+ | iCLIP: CITS, PureCLIP | GSE78030,GSM2064710 |
| ZNF674-AS1 (+)  ENST00000421685 | chrX:46546591-46546592;+  chrX:46545540-46545567;+  chrX:46546588-46546615;+  chrX:46546688-46546727;+  chrX:46546794-46546834;+  chrX:46546904-46546926;+  chrX:46546998-46547038;+  chrX:46547250-46547274;+  chrX:46547342-46547369;+  chrX:46547491-46547530;+  chrX:46547857-46547875;+  chrX:46547988-46548027;+  chrX:46548188-46548221;+ | iCLIP: PureCLIP  eCLIP | GSE78030,GSM2064710  ENCODE: ENCFF739LLZ, ENCFF086HIT |

(+) corresponds to transcripts from the positive strand

(-) corresponds to transcripts from the negative strand

**Table S2.** Functional and experimental details regarding the long noncoding RNAs used in this study.

| **LncRNA, RNA length, transcript ID** | **Function** | **Probing conditions and coverage** |
| --- | --- | --- |
| FTX (-)  2597 nucleotides ENST00000429124.5 | Positive regulator of XIST ^1^ | *NAI-N3 icSHAPE in HEK293* *^2^*  Chromatin *in vitro*: 0.6916  Chromatin *in vivo*: 0.6916  *DMS-MaPseq in HEK293T* ^3^  *In vivo*: 0.0111 |
| INE1 (+)  945 nucleotides ENST00000456273.1 | Continued expression is associated with prostate cancer, likely playing a role in autophagy regulation ^4^ | *NAI-N3 icSHAPE in HEK293* ^2^  Chromatin *in vitro*: 0.7704  Chromatin *in vivo* 0.7704  *DMS-MaPseq in HEK293T* ^3^  *In vivo*: 0.1143 |
| JPX (+)  2194 nucleotides ENST00000414209.5 | Activation of the XIST lncRNA  Aids in chromatin loop organization ^5–7^ | *NAI-N3 icSHAPE in HEK293* *^2^*  Chromatin *in vitro*: 0.6778  Chromatin *in vivo*: 0.6778  Cytoplasm *in vitro*: 0.1855 |
| MIR503HG (-)  646 nucleotides ENST00000457876.5 | Functions as the host gene of miR-503, mediating its expression  Affects cell proliferation, invasion, metastasis, apoptosis, and angiogenesis ^8–10^ | *NAI-N3 icSHAPE in HEK293* *^2^*  Chromatin *in vitro*; 0.0991  Chromatin *in vivo*: 0.0991  Nucleoplasm *in vitro*: 0.1084  *DMS, DIM-2P-seq in HEK293T* *^11^*  *In vivo*: 0.1130 |
| ZNF674-AS1 (+)  2078 nucleotides ENST00000421685.2 | Inhibits cell migration and proliferation  Inhibits proteins in glycolysis ^12,13^ | *NAI-N3 icSHAPE in HEK293* *^2^*  Cytoplasm *in vitro*: 0.0284  Nucleoplasm *in vitro*: 0.2820  *DMS, DMS-MaPseq in HEK293T* *^3^*  *In vivo*: 0.0823 |
| LINC00630 (+)  2101 nucleotides  ENST00000440496.5 | Promotes metastasis and the progression of non-small cell lung cancer  Associated with radioresistance of colorectal cancer ^14,15^ | *NAI-N3, icSHAPE in HEK293* ^2^  Chromatin *in vitro*: 0.1142  *DMS, DMS-MaPseq in HEK293T* ^2,3^  *In vivo*: 0.0609 |
| LINC00894 (+)  284 nucleotides  ENST00000455777.1 | Upregulated in lung cancer adenocarcinoma and thyroid cancer ^16,17^ | *NAI-N3, icSHAPE in HEK293* ^2^  Chromatin *in vitro*: 0.2042 |
| FIRRE (-)  2928 nucleotides  ENST00000427391.1 | Chromosomal organization  Positioning of X_i_ to the perinucleolar region via CCCTC binding factor (CTCF)  Pluripotency and adipogenesis ^18–20^ | *DMS, DMS-MaPseq in HEK293T* *^3^*  *In vivo*: 0.0888 |
| XIST RepA (-)  1260 nucleotides  ENST00000429829.5 | Transcriptional silencing of the inactive X-chromosome ^21,22^ | *NAI-N3, icSHAPE in HEK293* *^2^*  Chromatin *in vitro*: 1.0  Chromatin *in vivo*: 1.0  Cytoplasm *in vitro*: 1.0  Cytoplasm *in vivo*: 1.0  Nucleoplasm *in vitro*: 1.0  Nucleoplasm *in vivo*: 1.0  *NAI-N3, icSHAPE in HEK293T* *^23^*  *In vitro*: 1.0  *In vivo*: 1.0 |

**Abbreviations:** Human Embryonic Kidney 293 (*HEK293*), *In vivo* click selective 2-hydroxyl acylation and profiling experiment (*icSHAPE*), 2-(Azidomethyl)nicotinic acid imidazolide (*NAI-N3*), Dimethyl Sulfate Mutational Profiling Sequencing (*DMS-MaPseq*), DMS-induced mutations mapped by 2P-seq (*DIM-2P-seq)*

(+) corresponds to transcripts from the positive strand

(-) corresponds to transcripts from the negative strand

**Table S3.** iCLIP RBM15 crosslinking RNA binding sites for RBM15 with each lncRNA.

| **Binding Motifs** | **Stem-loop** | **Multiple stem-loops*** | **Single-stranded** | **Double-stranded and stem** |
| --- | --- | --- | --- | --- |
| **XIST A-repeats** | 5 | 6 | N/A | N/A |
| **FIRRE** | 4 | 1 | N/A | N/A |
| **INE1** | 3 | 3 | N/A | 2 |
| **MIR503HG** | 1 | N/A | 1 | 2 |
| **LINC00894** | 1 | 1 | N/A | N/A |
| **LINC00630** | 3 | 3 | N/A | N/A |
| **FTX** | 2 | N/A | N/A | 1 |
| **JPX** | 7 | 2 | N/A | 1 |
| **ZNF674-AS1** | 5 | 5 | N/A | 2 |
| **Total Count** | 31 | 21 | 1 | 8 |

Multiple stem-loops*: CLIP data extends more than one stem-loop

**Table S4.** Sequence logos display the predominant motifs, including CU, GA, and CA, commonly associated with stem-loop structures.

| **Stem-loop**  **Sequence Name** | **CU** | **GA** | **CA** |
| --- | --- | --- | --- |
| **XIST A-repeats** | 7, 9 | 24 | 2, 5, 15 |
| **FIRRE** | 3 | 6, 11, 14 | N/A |
| **INE1** | 2, 9, 19 | 4, 16 | 12, 21 |
| **MIR503HG** | 2, 11, 19, 28 | 24, 30 | 8, 14 |
| **LINC00894** | 18, 25, 38 | 29, 31, 44, 47, 51, 55 | 2, 5, 12, 15, 23, 41 |
| **FTX** | 6, 21, 28, 43 | N/A | 11, 23, 37, 40, 50 |
| **LINC00630** | 27, 37, 52, 54, 58 | 3, 8, 15, 19, 44 | 13, 35 |
| **JPX** | 5 | 3 | 7, 24 |
| **ZNF674-AS1** | 2, 21, 45, 48 | 9, 11, 25, 32, 39 | 27, 35, 50 |
